# The rotavirus VP5*/VP8* conformational transition permeabilizes membranes to Ca^2+^

**DOI:** 10.1101/2023.10.15.562449

**Authors:** Marilina De Sautu, Tobias Herrmann, Simon Jenni, Stephen C. Harrison

**Affiliations:** Department of Biological Chemistry and Molecular Pharmacology, Harvard Medical School, 250 Longwood Avenue, Boston, MA 02115, USA; Laboratory of Molecular Medicine, Boston Children’s Hospital, Boston, MA 02115, USA; Howard Hughes Medical Institute, Harvard Medical School, Boston, MA 02115, USA

## Abstract

Rotaviruses infect cells by delivering into the cytosol a transcriptionally active inner capsid particle (a "double-layer particle": DLP). Delivery is the function of a third, outer layer, which drives uptake from the cell surface into small vesicles from which the DLPs escape. In published work, we followed stages of rhesus rotavirus (RRV) entry by live-cell imaging and correlated them with structures from cryogenic electron microscopy and tomography (cryo-EM and cryo-ET). The virus appears to wrap itself in membrane, leading to complete engulfment and loss of Ca^2+^ from the vesicle produced by the wrapping. One of the outer-layer proteins, VP7, is a Ca^2+^-stabilized trimer; loss of Ca^2+^ releases both outer-layer proteins from the particle. The other outer-layer protein, VP4, activated by cleavage into VP8* and VP5*, is a trimer that undergoes a large-scale conformational rearrangement, reminiscent of the transition that viral fusion proteins undergo to penetrate a membrane. The rearrangement of VP5* thrusts a 250-residue, C-terminal segment of each of the three subunits outward, while allowing the protein to remain attached to the virus particle and to the cell being infected. We proposed that this segment inserts into the membrane of the target cell, enabling Ca^2+^ to cross. In the work reported here, we show the validity of key aspects of this proposed sequence. By cryo-EM studies of liposome-attached virions ("triple-layer particles": TLPs) and single-particle fluorescence imaging of liposome-attached TLPs, we confirm insertion of the VP4 C-terminal segment into the membrane and ensuing generation of a Ca^2+^ "leak". The results allow us to formulate a molecular description of early events in entry. We also discuss our observations in the context of other work on double-strand RNA virus entry.

## INTRODUCTION

Rotaviruses, like most other double-strand RNA (dsRNA) viruses, initiate infection of a cell by delivering an intact, inner capsid particle into the cytosol (Crawford et al., 2023). The two protein shells of this "double-layer particle" (DLP) enclose the 11 distinct segments of the dsRNA genome (Trask et al., 2012), each associated with an RNA-dependent RNA polymerase (RdRp) (Ding et al., 2019; Jenni et al., 2019) and an RNA cap-generating activity (Kumar et al., 2020; Ogden et al., 2014). On an infectious virion, a "triple-layer particle" (TLP), an outer protein shell surrounds the DLP (Figure 1A). Its role is to effect delivery of the DLP, which once liberated into the cytosol begins promptly to transcribe the dsRNA genome segments and to extrude them as capped mRNAs (Figure 1B).

**Figure 1.**
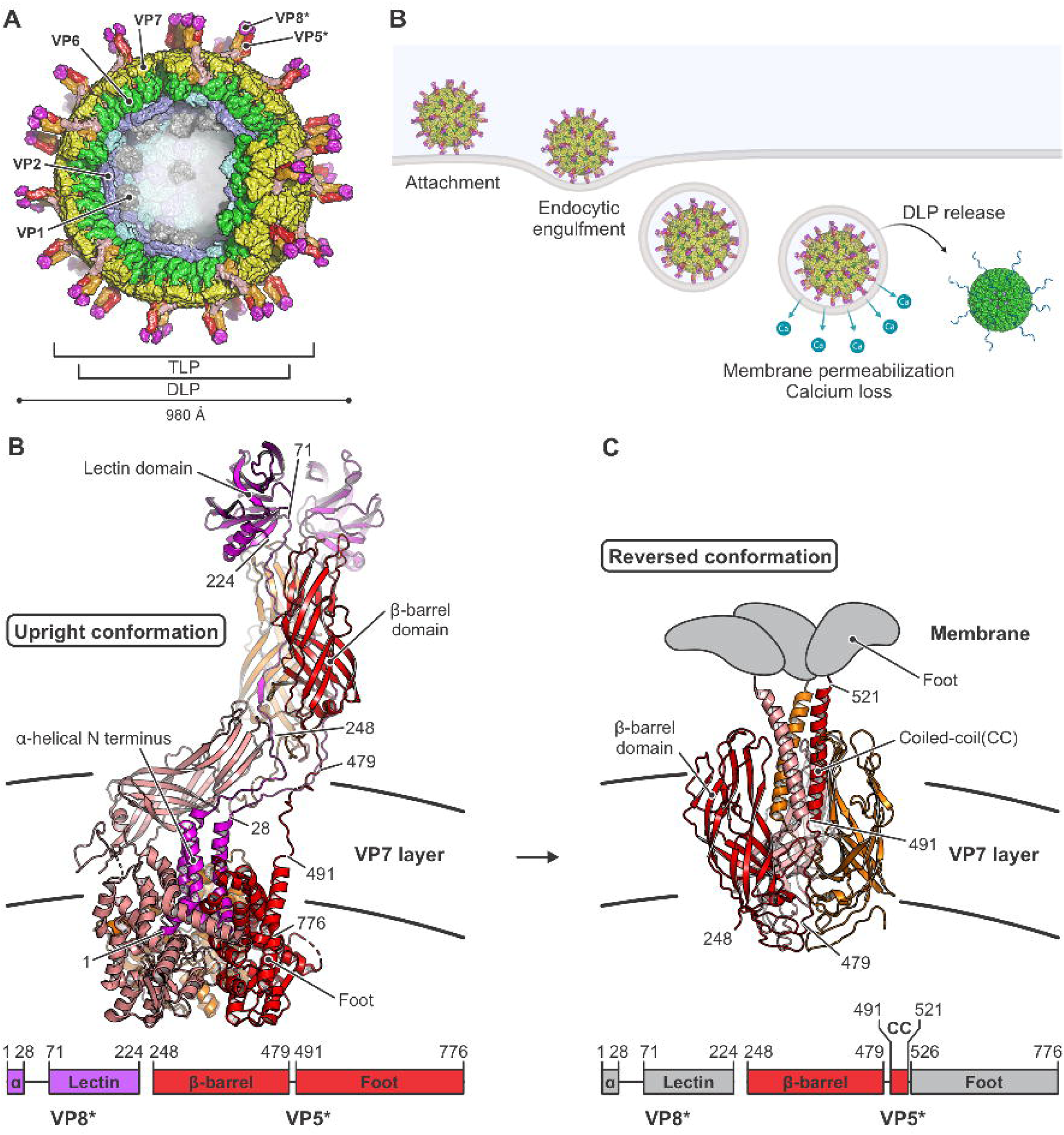
Rotavirus structure, entry, and conformational changes of its membrane-penetration protein VP4 (VP5*/VP8*). (A) Schematic illustration of an infectious rotavirus particle. Subunits are shown in surface representation and the particle is partially cut to allow inside view. The “double-layer” particle (DLP) is composed of VP2 (VP2A and VP2B, colored in blue and cyan, respectively) and VP6 (green). The outer layer of the “triple-layer” particle (TLP) also contains VP7 (yellow) and VP4 (VP5*/VP8* after proteolytic cleavage). VP5* is in red, orange and salmon; VP8*, in magenta; VP1, the capsid-bound RNA-dependent RNA polymerases (RdRp), in gray. The RNA cap-generating VP3, which has not been localized at a defined position in the virion, is not shown. (B) Schematic drawing of steps in rotavirus entry, summarizing observations from live-cell imaging (Abdelhakim et al., 2014; Salgado et al., 2018; Salgado et al., 2017) and biochemical and structural studies (Herrmann et al., 2021; Settembre et al., 2011). Entry requires trypsin-catalyzed cleavage of the VP4 spike protein into VP5* and VP8*. VP8* binding to glycolipid headgroups allows the TLP to attach to the surface of a host cell. This interaction, potentially in concert with membrane engagement of hydrophobic loops on VP5* (see panels C and D), allows the particle to enclose itself in an inward-budding vesicle. Subsequent steps include loss of Ca^2+^ from the vesicle interior, dissociation of VP7 and VP8*/VP5* from the DLP, free diffusion of the DLP in the cytosol, and initiation of RNA transcription. (C) Structure of the VP5*/VP8* spike in upright conformation (Herrmann et al., 2021), colored as in (A). The linear domain organization of the upright conformation is shown at the bottom. (D) Structure of the VP5* spike in reversed conformation (Herrmann et al., 2021), colored as in (A). The linear domain organization of the reversed conformation is shown at the bottom. Residues of the foot VP5* domains that were extruded from the VP7 layer cavity upon structural transition are shown schematically in gray.

Current evidence suggests that the two protein components of the outer layer, VP4 and VP7, have distinct functions in DLP delivery (Aoki et al., 2009; Herrmann et al., 2021; Kim et al., 2010; Settembre et al., 2011; Tihova et al., 2001). VP4, activated by proteolytic cleavage into VP8* and VP5*, is the principal membrane-interacting partner; VP7, held together by Ca^2+^ ions at its trimer interfaces, is a Ca^2+^-sensitive “clamp” that anchors VP4 onto the underlying DLP. For mammalian rotaviruses, cell attachment generally requires interaction of the VP8* moiety of VP4 with a glycolipid headgroup (Delorme et al., 2001; Dormitzer et al., 2002; Hu et al., 2012; Martinez et al., 2013; Ramani et al., 2013). For rhesus rotavirus (RRV) entering BSC-1 or SVG-A cells, uptake of the virus particle into a small vesicle does not require clathrin, dynamin, or related activities; the virus particle appears to wrap the plasma membrane around itself (Abdelhakim et al., 2014). Accompanying this step is a large-scale conformational rearrangement of the projecting VP4 trimer (cleaved to VP8* and VP5*) from an asymmetric, “upright” conformation (Figure 1C) to a threefold symmetric, “reversed” conformation (Figure 1D) (Herrmann et al., 2021). The transition takes place while the protein remains anchored on the virion surface by VP7. A major consequence of the reversal is projection outward of the ∼250 amino-acid residue “foot” segment of the VP5* polypeptide chain (Figure 1D). From our published structures and low-resolution cryogenic electron tomography (cryo-ET) (Herrmann et al., 2021), we concluded that the three copies of this foot segment probably insert into the plasma membrane of the target cell, permeabilizing the membrane to Ca^2+^ ions. Subsequent steps are loss of the Ca^2+^ ions that stabilize VP7 trimers and hence hold together the outer layer of the particle, dissociation of the outer layer, and escape of the DLP into the cytosol (Salgado et al., 2018) (Figure 1B).

We report here that we can recapitulate with liposomes early steps in the entry process – attachment to glycolipid headgroups and insertion of the projected foot domain into the lipid bilayer, as visualized by cryogenic electron microscopy (cryo-EM). We show by single-particle fluorescence imaging that this same interaction leads to passage of Ca^2+^ across the liposome membrane, but not to further disruption of the liposome. We relate these observations to the entry steps observed by live-cell imaging.

## RESULTS

### TLP permeabilization of liposomes to Ca^2+^ ions

Loss of Ca^2+^ from TLPs is an early step following engulfment at the cell surface (Salgado et al., 2018). We tested whether an interaction of virions with a membrane could lead to Ca^2+^ permeability, without contribution of potential cellular factors, using liposomes containing, in their lumen, a soluble, Ca^2+^-sensitive fluorophore (Fluo-4) and, in their bilayer, a fluorescently labelled lipid (Cy5-DOPE) for visualization, as well as biotinylated lipids for attachment to an avidin-coated coverslip. We prepared the liposomes in the absence of Ca^2+^ and used a lipid composition similar to that of mammalian-cell plasma membranes supplemented with 2.5% GD1a, a validated rotavirus attachment factor (Delorme et al., 2001; Dormitzer et al., 2002; Hu et al., 2012; Martinez et al., 2013; Ramani et al., 2013). In the absence of Ca^2+^, we established the background fluorescence of the incorporated Fluo-4 dye. We incubated the liposomes with fluorescently labeled TLPs (Atto 565) in Ca^2+^-containing buffer, then deposited the sample onto a coated coverslip and recorded signal at appropriate wavelengths by total internal reflection fluorescence (TIRF) microscopy (Figure 2A and B, Figures S1 and S2). Thus, the intensities of the diffraction-limited spots in each of the three recorded channels reported, respectively, the presence of liposomes, the presence of Ca^2+^-liganded Fluo-4 (and hence permeabilization of the liposome to Ca^2+^), and the attachment of TLPs. In a parallel control experiment, we used ionomycin, a Ca^2+^ ionophore, to permeabilize similarly prepared liposomes.

**Figure 2.**
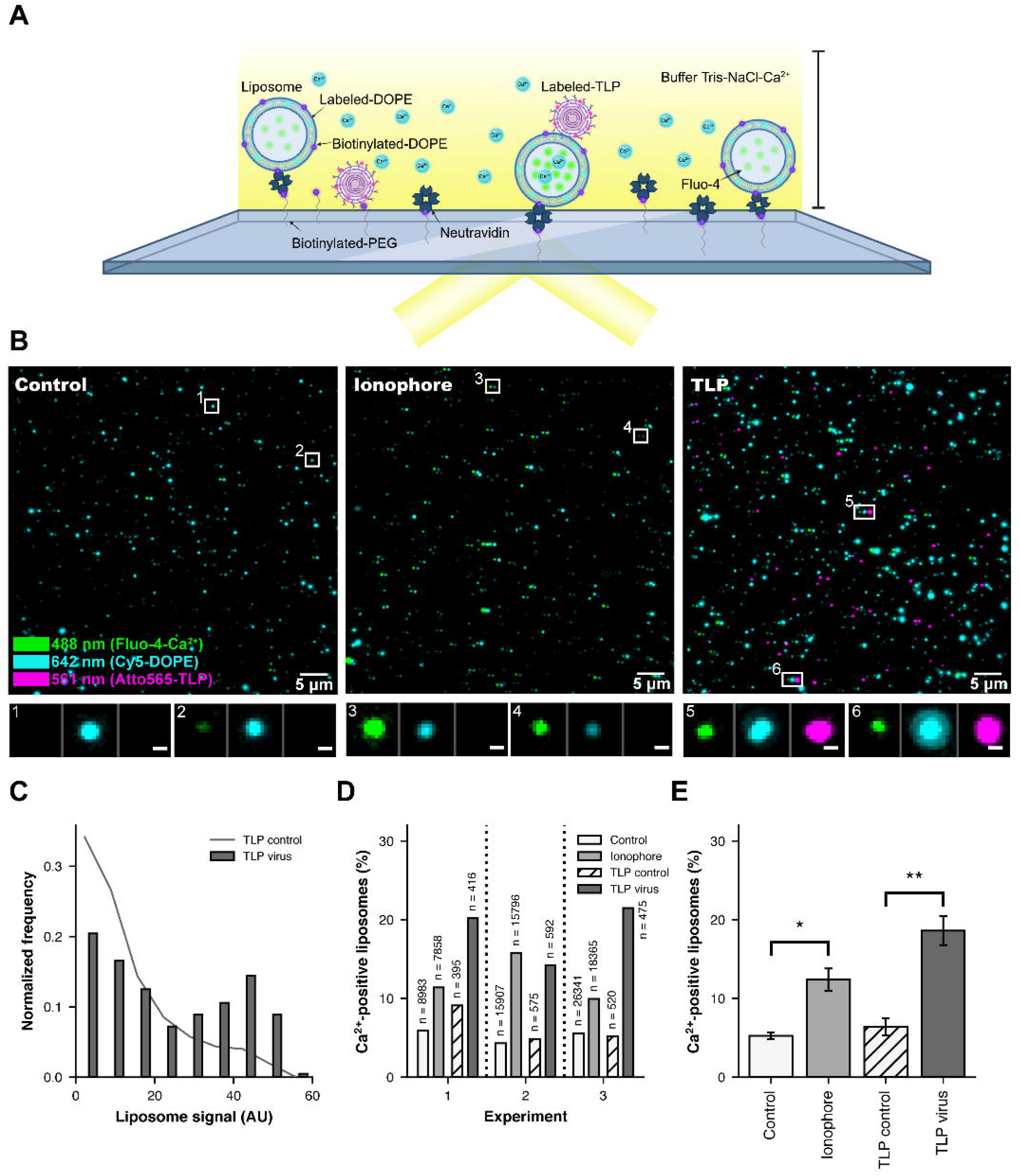
Interaction of rotaviruses with liposomes makes membranes Ca^2+^ permeable. (A) Schematic overview of the TIRF experimental setup. (B) Representative fluorescence images, showing a field of view for the control (left, liposomes only), ionophore (center, liposomes incubated with ionomycin) and TLP (right, liposomes incubated with TLPs) samples. The 640 nm channel (Cy5-DOPE, liposome signal) is in cyan and shifted 7 pixels to the right in all images to visualize colocalization; the 488 channel (Fluo-4, Ca^2+^ sensor signal) is in green; the 561 channel (Atto 565 NHS ester dye, virus signal) is in magenta and shifted 14 pixels to the right in the TLP sample. The scale bar corresponds to 5 μm. Representative examples of liposomes are boxed (1–6) and displayed beneath the images at higher magnification (scale bar = 0.5 μm) with the three channels shown next to each other. (C) Normalized frequency of the liposome signal from the TLP sample (experiment 1). The dark grey line shows the distribution for TLP control liposomes (liposomes that did not colocalize with a virus particle in the TLP sample) (n = 28,985). The dark gray bars show the distribution for the liposomes that colocalized with a virus (n = 415). (D) Ca^2+^-positive liposomes (%) for the control sample (white bars), the ionophore sample (light grey bars), the TLP control (striped white), and TLP virus (dark grey) samples from experiments 1, 2 and 3, respectively. "Ca^2+^-positive liposomes" means percent of the liposomes that had a Ca^2+^ signal with an intensity greater than 1.5 standard deviations from the mean (see red dashed lines in Figure S2A and S2C for the cutoffs). N = total number of liposomes in each condition. (E) Ca^2+^-positive liposomes (%) for the control sample (white bars), the ionophore sample (light grey bars), the TLP control (striped white), and TLP virus (dark grey) samples from the three independent experiments (Figures S1 and S2). Error bars are the standard error calculated from the mean of the independent experiments. Statistical analysis used a t-test between samples. * = p<0.05; ** = p<0.01.

The distribution of intensities for the lipid marker (Cy5-DOPE) showed that the liposomes varied in size over an extended range but that this range was very similar for all samples and experiments (Figure S1B). The larger liposomes were more likely to have a colocalized TLP, suggesting that the particles had associated preferentially with the larger liposomes, as might be expected from their greater cross section (Figure 2C). Some control liposomes, incubated with neither ionomycin nor TLP, showed Ca^2+^ positive signal, corresponding to random lesions, which varied somewhat from sample to sample (Figure 2B and 2D and Figure S2). Addition of TLPs yielded a markedly enhanced frequency of Ca^2+^ penetration (Figure 2D), at levels comparable to the enhancement by ionomycin (compare Figures 2D and 2E; Figure S2). For the overall effect shown in Figure 2E, we calculated the mean number of Ca^2+^-positive liposomes from three independent experiments (Figure S1 and S2) by counting the number of particles that showed a Ca^2+^ signal with an intensity above 1.5 standard deviations from the mean. Despite permeabilization to Ca^2+^, the liposomes remained impermeable to Fluo-4. That is, the interaction with TLPs did not, under these conditions, disrupt the integrity of the liposome. We conclude that binding of an RRV TLP to a membrane containing an appropriate sialic-acid receptor permeabilizes of the liposome membrane to Ca^2+^ ions, without otherwise rupturing the lipid bilayer.

### Cryo-EM analysis of TLPs with liposomes

Is the Ca^2+^ permeabilization detected by our fluorescence measurements a result of the VP5*/VP8* upright-to-reversed transition characterized previously? We visualized by cryo-EM the interaction of TLPs with lipid-bilayer membranes (Figure 3A), using liposomes prepared as in the experiments above, but without Fluo-4 and without the membrane label. We used recoated particles (rcTLPs), because of previous observations that our recoated TLPs were more infectious than virions isolated from infected cells, which can lose VP4 during purification, and also because we had found that VP4 on rcTLPs appeared to undergo the transition from upright to reversed more readily (Herrmann et al., 2021). We recorded four cryo-EM data sets on a Thermo Fisher Titan Krios operated at 300 kV and equipped with a Gatan Bioquantum energy filter and K3 detector (Table S1).

**Figure 3.**
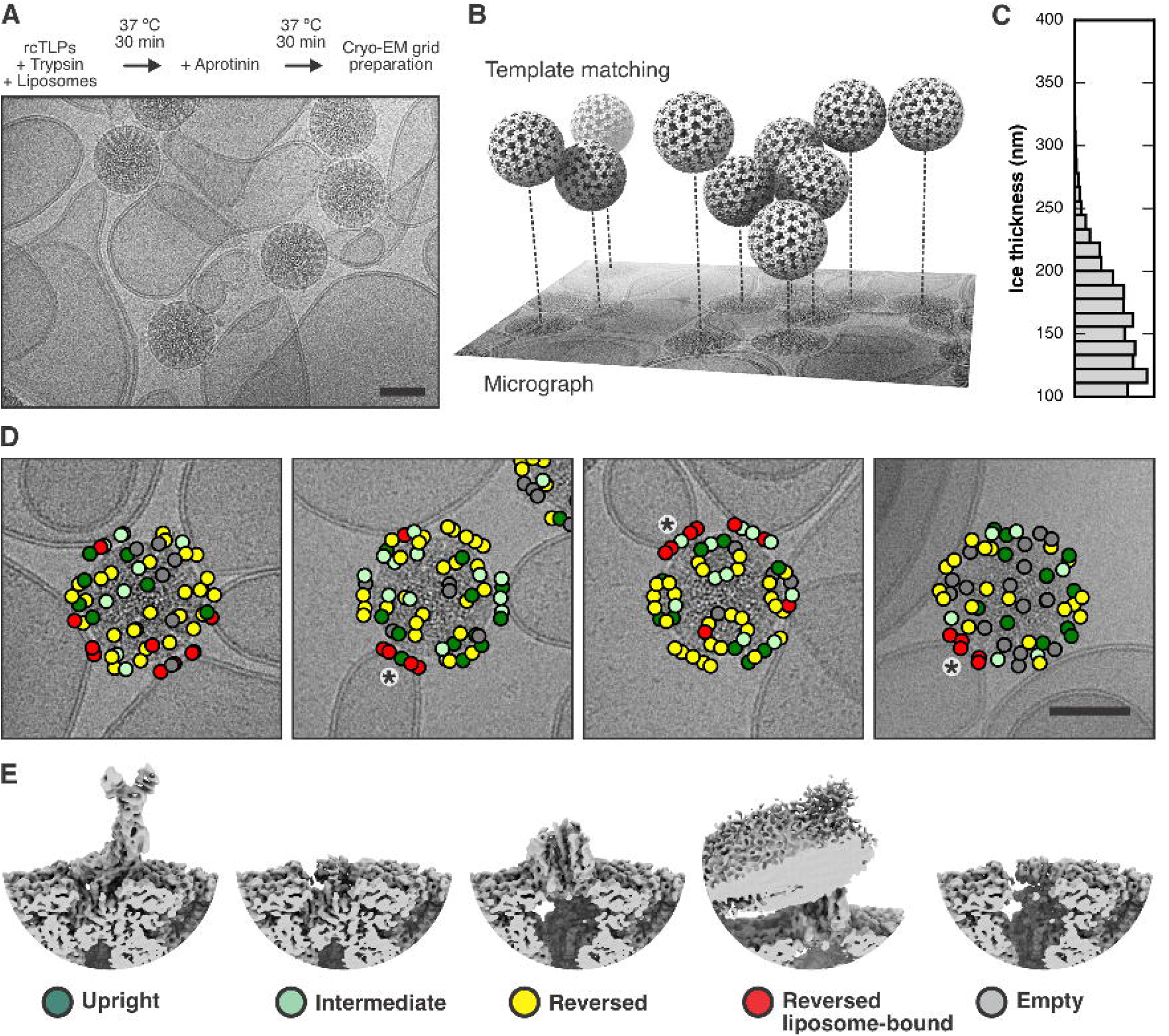
Cryo-EM analysis of rotavirus in the presence of liposomes. (A) Sample preparation scheme and representative micrograph showing rcTLP particles bound to liposomes. The scale bar corresponds 50 nm. (B) Illustration of how template matching was used to locate virions in the micrographs and estimate their z position based on per-particle defocus fitting. (C) Histogram of estimated ice thickness of the cryo-EM samples. For each micrograph that contained at least two particles, we calculated the largest z difference and added 100 nm (approximately two times the particle radius). (D) Mapping of spike positions and their class assignment onto the micrographs of liposome-bound viruses, color-coded as ©(E). The asterisks indicate a cluster of VP5* trimers around the five-fold axis of the virus. The scale bar corresponds 5©m. (E) Representative reconstructions of the different spike conformations after subparticle classification.

We found the position of viral particles on our micrographs by template matching from cisTEM (Grant et al., 2018; Lucas et al., 2021). By refining the local defocus for each particle, we obtained its relative *z* position within the vitreous ice (Figure 3B). Figure 3C shows a histogram of the largest *z* difference plus 100 nm (approximately two times the radius of a virion), for micrographs from which we picked at least two virus particles. From this distribution, we estimated that the ice thickness of our samples varied between 100 and 250 nm.

Images in Figure 3D show that many particles appeared to have extended contacts with liposomes, but many of these were with liposomes of radius ≥300 nm and hence substantially flattened in the ice. As a consequence of flattening, the bilayer would have been forced to curve away from the particle, not just in the plane of the ice but in the directions normal to it, probably peeling the membrane away from the particle during blotting (Figure S3). Moreover, as the suspension thinned, attached particles would have migrated to the periphery of the flattening liposome in order to remain in solution, distorting the liposome in the process. Thus, the invaginations seen at the more extended contacts appear to be largely within the plane of the image and of limited extent above and below it, and our images probably do not represent the full extent of contact achieved in the preparation. We can detect no evidence of liposome disruption when TLPs bind or when the interacting VP5* has transitioned from upright to reversed conformation.

An icosahedral reconstruction carried out after merging virus images from all four datasets resulted in a density map with an overall resolution of the viral shell of 2.43 Å (Figure S4 and Table S1). As described in Methods and in Figures S5 and S6, we then used subparticle classification (Figures 3D and E), starting with a total of 4,550,760 masked VP4 subparticles, to separate upright, intermediate, reversed, and empty VP5*/VP8* spikes positions (classifications #1 and #2, Figure S7 and Table S2) and to distinguish membrane-engaged from non-engaged spikes (classifications #3 and #4, Figure S8 and Table S3). Two relatively heterogenous classes showed membrane-engaged reversed spikes, with apparently variable angles of spike contact (Figure S8A, classification #3). A further classification step then yielded two maps from 56,593 and 70,018 subparticles (classes 5 and 6), respectively, with relatively well-defined lipid-bilayer density (Figure S8B, classification #4). Even within these classes, variability of curvature at positions at which the bilayer bent away from the particle blurred the membrane profile around the circumference of the contact. Local resolution estimates for classes 5 and 6 showed a well-resolved, virion-bound VP5* β-barrel trimer with its central coiled-coil domain and a poorly resolved lipid bilayer (Figure S9).

The class 6 reconstruction has pronounced lipid bilayer density on one side of the reversed VP5* and somewhat fuzzier density on the side opposite (Figure 4). The angle between the curving bilayer and the VP5* axis shows that this class represents a reversed spike near the edge of the contact. The side with a well-defined profile faces the nearest fivefold axis (Figure 4A and B). A fivefold axis may be close to the center of many of these membrane contacts, because the five surrounding spikes can offer a compromise between a minimally distorted membrane region and a set of membrane contacts stable enough to resist the forces during thinning of the aqueous film, as described above (see diagram in Figure S3). We verified that many of the subparticles contributing to this class indeed came from reversed VP5* spikes related by a five-fold axes, by mapping the class assignment onto the original micrographs (Figure 3D and E). The coiled-coil at the C-terminal end of the VP5* β-barrel domains extends just to the inner lipid headgroup layer, and diffuse density extends beyond it into the bilayer profile, but peeling away of the bilayer leaves nearly all the engaged spikes at the periphery of the contact, and the resolution leaves uncertain the molecular features of the likely membrane-penetrating elements (Figure 4C and D).

**Figure 4.**
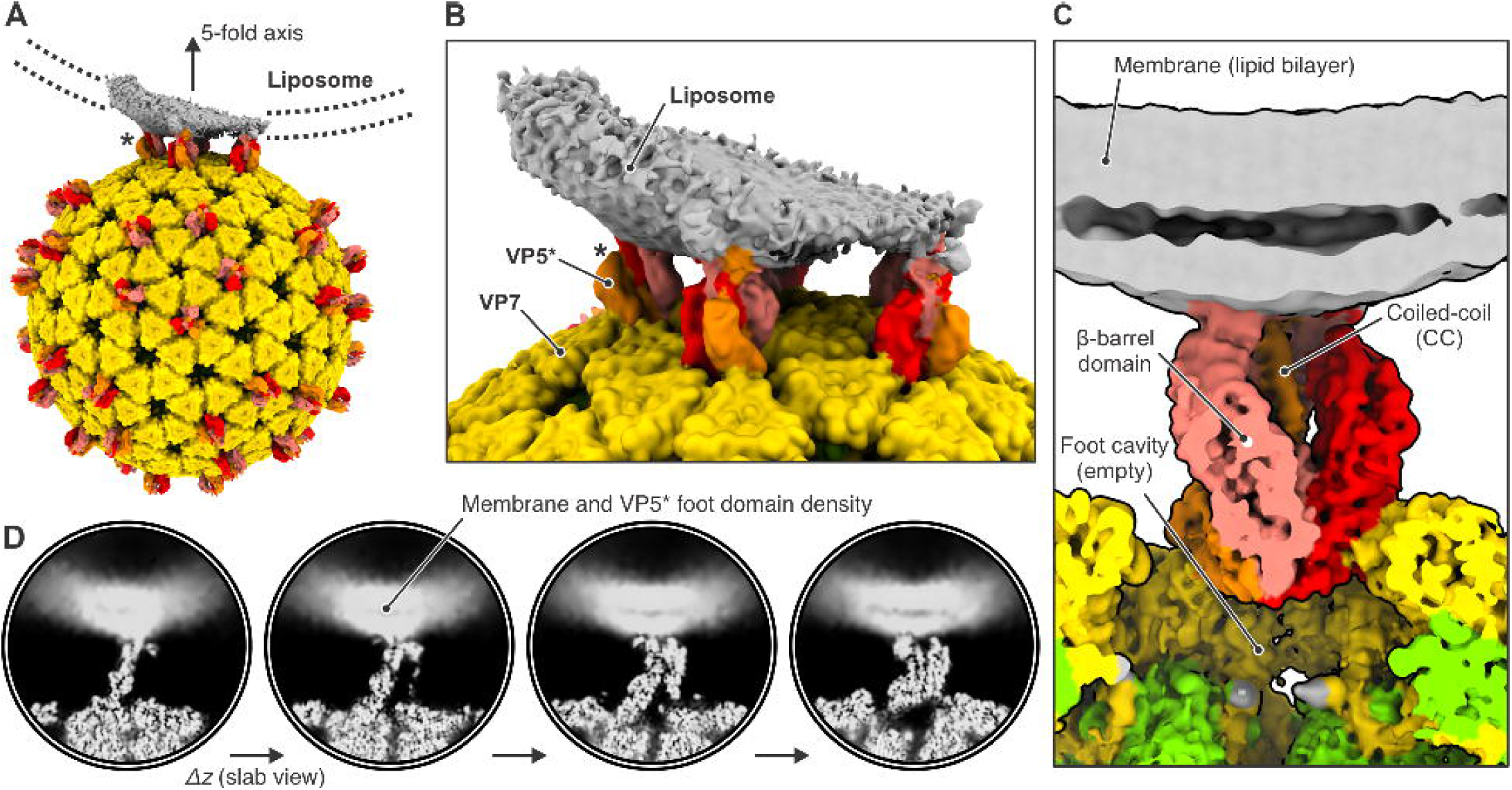
Cryo-EM reconstruction of membrane-bound VP5* in the reversed conformation. (A) Reconstruction of class 6 (Figure S8B) in context of the full virus. The map was low-pass filtered at 8.0 Å resolution. The outer-shell VP7 is colored yellow; VP5* is colored red, orange, and salmon; membrane density is gray. The asterisk denotes the VP5* trimer on which the subparticle classification was focused. The direction of one of the icosahedral five-fold axis is indicated. (B) View around the five-fold axis showing multiple VP5* trimers interacting with liposome. The asterisk denotes the VP5* trimer on which the subparticle classification was focused. (C) Close-up view of the membrane-bound VP5* trimer. The map is partially cut. (D) Slab views of the reconstruction showing diffuse lipid bilayer and membrane-embedded VP5* foot domain density.

The reversed-spike structure (PDB-ID 6WXG) (Herrmann et al., 2021) can be docked with confidence into the observed subparticle density (Figure 5). The tips of the VP5* β barrel are hydrophobic loops (Kim et al., 2010). Two of the three tips contact the liposome bilayer, while its curvature from flattening has probably eliminated a contact from the third. Mutations that reduce the hydrophobicity of these loops inhibit viral entry (Kim et al., 2010), and interpretation of tomographic reconstructions of TLPs entering at the thin edges of BSC-1 cells has suggested that these loops interact with membrane directly (Herrmann et al., 2021). Our current reconstruction is consistent with this interpretation and suggests that the interaction might already be present in a proposed intermediate state in which the β-barrel domains have shifted into the threefold arrangement found in the reversed structure, but the foot domains have not yet begun to project outward from their positions in the pre-attachment TLP (Herrmann et al., 2021).

**Figure 5.**
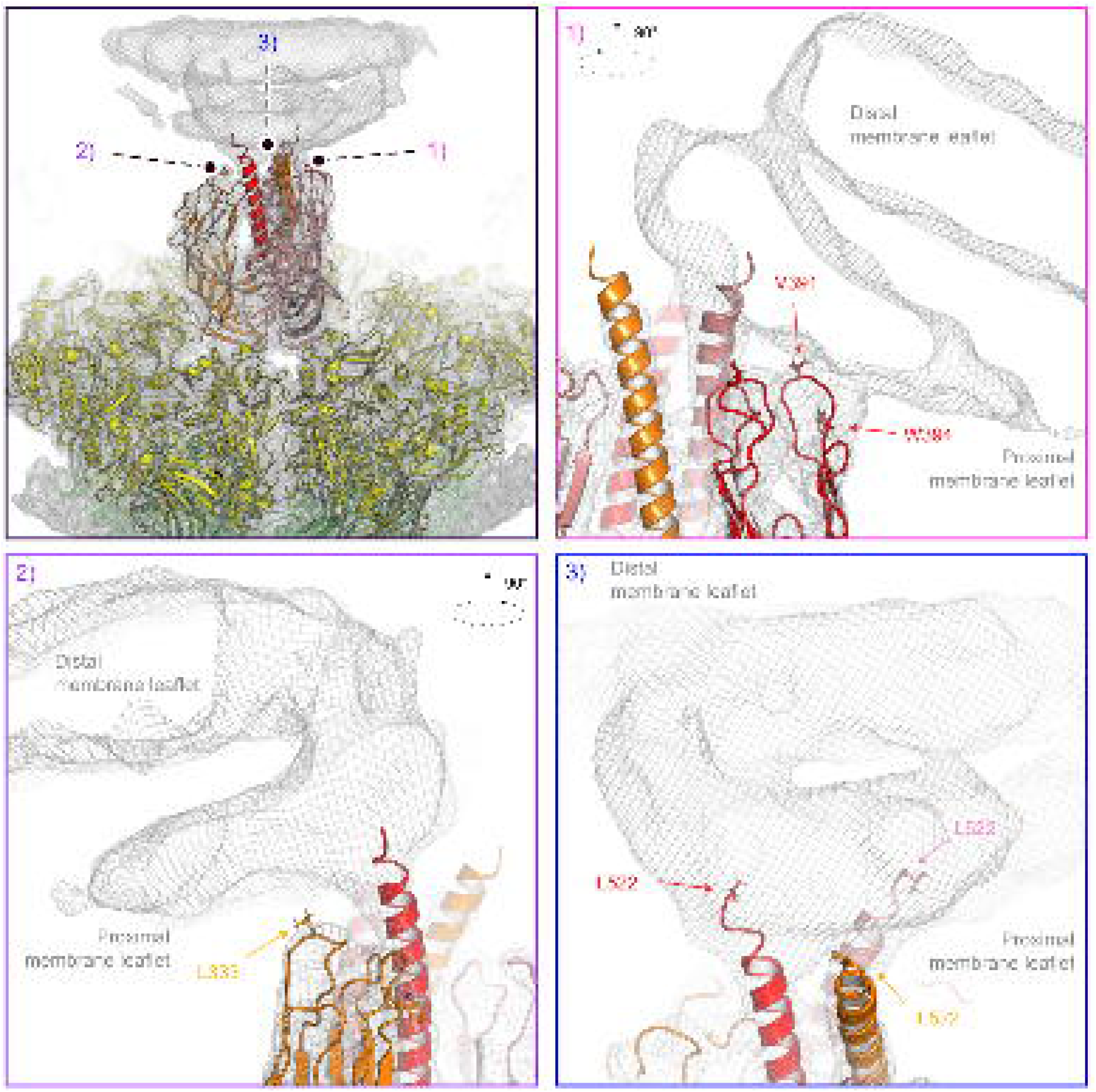
Interactions between the reversed VP5* trimer and the membrane. (A) Placement of the rotavirus-bound VP5* trimer structure in reversed conformation (PDB-ID 6WXG) (Herrmann et al., 2021) into the density map of the class 6 reconstruction (Figure 4). Density is shown as gray mesh, docked viral protein subunits are shown in ribbon representation and colored yellow (VP7); red, orange and salmon (VP5*). (B) Close-up view of the hydrophobic loops at the tip of one VP5* β-barrel that interact with the proximal membrane leaflet. (C) Close-up view of the hydrophobic loops at the tip of a second VP5* β-barrel that interact with the proximal membrane leaflet. (D) Close-up view of trimeric VP5* coiled-coil extending into the membrane.

## DISCUSSION

We have shown in published work (Herrmann et al., 2021), which combined cryo-EM single-particle analysis with cryo-ET of engulfed virions entering cells, that VP5* on the surface of an attached TLP can undergo a transition from an upright to a reversed conformation without dissociating from the particle, causing its C-terminal foot domain to project into the apposed membrane (Figure 1B and C). DLPs recoated with VP7 and a foot-locked VP4 mutant that cannot undergo the full reversal have substantially lower infectivity than do those recoated with VP7 and wild-type VP4 (Herrmann et al., 2021). Projection of the foot into the surrounding vesicle is therefore essential for infection.

We have now shown, by analysis of individual spike positions on cryo-EM images of virions attached to liposomes, that VP5* trimers, in reversed conformation, interact with the lipid bilayer, recapitulating the interaction inferred from icosahedral averaging of much lower resolution cryo-ET images (Figures 4 and 5). The reconstruction shows that the central coiled-coil domain protrudes into the lipid bilayer, although flattening and distortion of the liposomes have prevented us from resolving protein structure within the bilayer. Under essentially identical conditions, TLPs that attach to similar liposomes allow Ca^2+^ ions to pass into the liposomes without destroying their integrity --that is, the permeabilization step detected by live-cell imaging (Abdelhakim et al., 2014). In the experiments here, Ca^2+^ enters a liposome; during viral entry, it leaves the vesicle surrounding the TLP, driven by the concentration gradient from vesicle to cytosol.

Combining these results with those from earlier work, we can summarize as follows our current understanding of the molecular events accompanying RRV entry.

1. Initial attachment to sialoglycolipids on the surface of a cell is through VP8* at the apex of the VP8*/VP5* spike in its upright conformation (Abdelhakim et al., 2014).
2. Engulfment (disappearance from the cell surface into the cytoplasm) is complete within a few minutes for most of the attached particles. DLPs recoated with VP4 mutated in the hydrophobic loops of VP5* move from the cell surface into the cell interior, but fail to release into the cytosol (Abdelhakim et al., 2014) and fail to infect (Kim et al., 2010). That is, engagement of the hydrophobic loops with the membrane bilayer appears not to be necessary for particle uptake but essential for later steps. Complete enclosure from an essentially flat lipid bilayer would require that interaction of the virion with the membrane liberate a total free energy of about 200 kcal/mol of vesicle (but substantially more for 30-40 nm liposomes) (Helfrich, 1973). The K_d_ of RRV VP8* for sialic acid is 1.2 mM; a rough calculation of the free energy derived from binding, taking as a local concentration of sialic acid headgroups as the number of gangliosides within 50 Å of a VP8* site and the ganglioside concentration in the membrane as 1% of the phospholipids, suggests that the free energy of binding could indeed generate enough work (≥200 kcal/mol of particle) to drive complete wrapping of the membrane, assuming high VP4 occupancy. In practice, gangliosides and other sphingolipids tend to cluster transiently into so-called "lipid rafts", so that the local concentration might be considerably higher than the cell-surface average.
3. VP4 on the surface of a virus particle can transition spontaneously into its reversed conformation. Why do the engulfed hydrophobic-loop mutants fail to release and infect? A possible explanation is that when the hydrophobic loops cannot engage the membrane, the projected foot fails to insert, and the reversal is futile. Formation of the central coiled-coil presumably drives foot extrusion, but loop engagement (or some attachment stronger than VP8* binding to ganglioside headgroups) may be necessary to resist the kinetic barrier to membrane insertion. The explanation is consistent with the published *in vitro* observation that release of cleaved VP4 (i.e., VP8*/VP5*) from TLPs (by chelating Ca^2+^) in the presence of liposomes leads to liposome association of VP5* trimers, but not if liposomes are added later (Trask et al., 2010).
4. What initiates the reversal? The interface between the VP8* lectin domain and VP5* is modest, and that domain alone would probably dissociate readily from its VP5* partner. The VP8* N-terminal, coiled-coil anchor in the foot will prevent full dissociation from the virion. However, even if the lectin domain fluctuates on and off that fluctuation will expose the VP5* hydrophobic loops, which can engage a membrane when VP8* has attached to the lipid headgroups. This picture is consistent with our suggestion, above, that hydrophobic loop engagement is essential for foot insertion. It is also consistent with our finding for a human rotavirus vaccine-strain candidate, that mutations during passage, including one at the VP8*/VP5* interface, apparently stabilize the upright conformer (Jenni et al., 2022).
5. Foot insertion allows Ca^2+^ to cross the vesicle membrane. If reversal and membrane insertion occur before complete engulfment, Ca^2+^ will pass into the cytosol from the extracellular medium. It is indeed possible that Ca^2+^ signaling by this mechanism ordinarily accompanies rotavirus entry. Foot insertion within a closed vesicle will quickly deplete the contents of Ca^2+^, entraining outer-layer dissociation and subsequent events, which must include further disruption or perforation of the vesicle membrane.

Orbiviruses, as represented by bluetongue virus, appear to have a similar mechanism for membrane insertion and endosome penetration (Xia et al., 2021; Zhang et al., 2016). Bluetongue virus enters through low-pH compartments, and the signal for conformational change is proton binding, probably by a cluster of histidines, rather than Ca^2+^ loss. The bluetongue virus subunit VP5 (the functional analog of rotavirus VP5*, but with a quite different, trimeric structure) undergoes, when the pH drops to 6 or below, a stepwise "unfurling" of an antiparallel coiled-coil, thereby forming a ∼190 Å long stalk, at least part of which is a six-helix bundle (Xia et al., 2021). Residues in the three loops at the distal tip of this bundle appear to insert into liposomes when the latter are mixed with virus particles *in vitro*. The mechanism for release of the core particle (the DLP equivalent) from an endosome may involve a wrapping of endosomal membrane around the low-pH altered virion.

Orthoreoviruses initiate penetration by releasing a pore-forming, myristoylated peptide (μ1N), generated by autolytic cleavage of the outer-shell protein, μ1 (Agosto et al., 2006; Zhang et al., 2009). The pores formed *in vitro* in red blood cells or liposomes are about 50 Å in diameter, but they could in principle be larger when μ1N release occurs within an endocytic vesicle. A molecular mechanism for the ultimate release of a core particle into the cytosol is still uncertain.

The rotavirus infectious cycle includes another membrane-crossing step, when a newly assembled DLP, together with VP4, acquires its VP7 complement in the endoplasmic reticulum (ER). VP7, a glycoprotein, folds in the ER; inhibition of its synthesis or of its proper folding leads to accumulation in the ER of enveloped DLPs (Altenburg et al., 1980; Poruchynsky and Atkinson, 1991; Soler et al., 1982). Recent cryo-ET studies under conditions of VP7 inhibition have captured VP4-decorated DLPs budding into the ER (Shah et al., 2023), presumably mediated by the virally encoded ER receptor, NSP4 (formerly NS28) (Bergmann et al., 1989). Whether the resulting enveloped particles are an on-pathway intermediate or a dead-end consequence of VP7 inhibition remains to be established. It is possible that VP7 trimers participate in productive budding, rather than displacing an already complete bilayer from around a budded particle. It likewise remains possible that VP7 participates directly in the membrane disruption required for complete release of the DLP during viral entry -- the step for which we now seek a molecular picture.

## METHODS

### Cells, plasmids and constructs

MA104 cells (ATCC, type: MA104 clone 1, ATCC CRL2378.1, cat. # ATCC CRL2378.1) were grown in M199 medium (Thermo Fisher Scientific) supplemented with 25 mM 4-(2-hydroxyethyl)-1-piperazineethanesulfonic acid (HEPES) and 10% Hi-FBS (Thermo Fisher Scientific). BSC-1 cells (ATTC) were grown in DMEM (Thermo Fisher Scientific) supplemented with 10% Hi-FBS (Thermo Fisher Scientific). For VP4 expression, we cloned its full-length genomic sequence from serotype P5B[3], NCBI:txid444185) into a pFastbac expression vector.

For VP7 expression, we cloned its full-length genomic sequence (G3 serotype, NCBI:txid444185) into a pFastbac expression vector and transformed *E. coli* DH10α cells. For bacmid generation, VP4 and VP7 vectors were transformed into DH10-Bac cells (ThremoFisher Scientific) and plated onto LB-agar plates supplemented with 50 μg/ml kanamycin, 7 μg/ml gentamycin, 10 μg/ml tetracycline, 100 μg/ml blue-gal and 40 μg/ml β-D-1-thiogalactopyranoside (IPTG).

### Reagent preparation

#### Purification of rotavirus DLPs and TLPs

We grew MA104 cells to confluency in 850 cm^2^ roller bottles in M199 medium supplemented with 10% FBS and 25 mM HEPES. Cells were infected with rhesus rotavirus (RRV, strain G3P5B[3]) at a multiplicity of infection (MOI) of 0.1. After incubation at 37 °C for 24 h, the medium containing cells and debris was collected and frozen at -80 °C for storage. To purify DLPs, we uncoated TLPs by adding EDTA pH 8.0 to a final concentration of 5 mM. After thawing, we pelleted cells and debris by centrifugation in a Beckman Coulter rotor JS 4.2 at 3,200 rpm for 30 min at 4 °C. Supernatant was then removed and the pellet resuspended in 1 ml ice cold TNE buffer (20 mM Tris, 100 mM NaCl, 1 mM EDTA, pH 8.0) for DLP purification. We concentrated viral particles from the supernatant by ultracentrifugation in a Beckman Coulter rotor Ti-45 rotor at 44,000 rpm. The supernatant was then discarded and the pellet resuspended in 1–3 ml ice cold TNE and combined with the cell-debris fraction from the first centrifugation. We transferred the resulting suspension to a 15 ml conical tube and added TNE to a final volume of 4 ml. We added 4 ml of 1,1,2-trichloro-1,2,2-trifluoroethane (Freon 113) and inverted the tube 5–10 times. The suspension was centrifuged for 2 min in a Beckman Coulter rotor SX4750 at 1,000 rpm at 4 °C. We removed the upper aqueous phase, transferred it to a new clean 15 ml conical tube and repeated the freon extraction two more times. The resulting virus solution was layered on top of a 34.7–60.0% (w/v) CsCl gradient (1.26–1.45 g/ml). CsCl solutions were prepared in either TNE (DLP purification) or TNC (TLP purification). Gradients were spun at 4 °C in a Beckman Coulter rotor SW 60 at 55,000 rpm for 2.5 h. Bands for TLPs and DLPs were harvested by side puncture. TLPs and DLPs were dialyzed against 2 L of TNC or TNE overnight. Viral particles were concentrated by pelleting at 4 °C in a Beckman Coulter rotor SW 60 at 55,000 rpm for 1 h. Supernatants were removed with a 1 ml pipette and pellets resuspended in 100–200 μl of remaining buffer. Concentrations of DLPs were determined with a nanodrop measuring the absorption at 260 nm (extinction coefficient = 4.7028 g^-1^cm^-1^). TLP concentrations were measured by densitometry of VP6 bands on Coomassie stained SDS-PAGE gels against DLP standards ranging from 0.1 to 1.0 mg/ml.

#### Purification of recombinant VP7

We initially obtained baculovirus carrying the genetic information of full-length rhesus rotavirus VP7 (G3 serotype) from bacmid transfected Sf9 cells (Thermo Fisher Scientific). Sf9 cells were then inoculated with baculovirus and passaged in the same cells three times, with an incubation time of 72 h for each passage. Eight flasks of 500 ml Sf9 cells with approximately 2 million cells per ml were then infected with 13 ml of passaged virus stock solution and incubated for 72 h. Cells were harvested at 4 °C by centrifugation in Beckman Coulter rotor J 4.2 at 3500 rpm for 30 min. Supernatant was transferred to a clean 4 L beaker and benzamidine and sodium azide were added to final concentrations of 1 mM and 0.01% (w/v), respectively. 50 ml of a concanavalin A Sepharose (ConA) resin was added to the supernatant and stirred at 200 rpm overnight. Using gravity flow, the resin was packed into a column and washed with five column volumes (CV) of TNC buffer. Protein was eluted with five column volumes of TNC buffer containing 0.6 M α-methyl mannose. Using a peristaltic pump, we loaded the eluate onto 11 ml protein A resin with immobilized antibody m159 (Greenberg et al., 1983) (10 mg per ml of protein A resin), specific for trimeric VP7, and equilibrated in buffer A (20 mM Tris, 50 mM NaCl, 0.1 mM CaCl_2_, pH 8.0). We washed bound VP7 protein with five CV of buffer, and eluted with five CV of buffer B (20 mM Tris, 50 mM NaCl, 1 mM EDTA, pH 8.0). Buffer was exchanged using a G25 PD10 column equilibrated in 0.1HNC (2 mM HEPES, 10 mM NaCl, 0.1 mM CaCl_2_) spiked with 0.1 mM PMSF. Protein was divided into aliquots, which were frozen in liquid nitrogen and stored at -80 °C.

#### Purification of recombinant VP4

We harvested baculovirus carrying the genetic information of full-length VP4 (P5B[3] serotype) from bacmid transfected Sf9 cells. Sf9 cells were then inoculated with baculovirus and passaged in the same cells three times, with an incubation time of 72 h for each passage. Four flasks of 500 ml Sf9 cells grown in suspension to a density of approximately 2 million cells per ml were then infected with 13 ml of passaged virus stock solutions and incubated for 72 h. Cells were harvested at 4 °C by centrifugation in Beckman Coulter rotor J 4.2 at 3,500 rpm for 30 min. We discarded the supernatant, resuspended the cell pellet in 100 ml lysis buffer (75 mM Tris, 100 mM NaCl, 5 mM EDTA, 7.5% (v/v) glycerol, 1 mM PMSF, 1 mg/ml aprotinin, 1 mg/ml pepstatin, 1 mg/ml leupeptin), and froze the sample in liquid nitrogen for storage at -80 °C. After thawing of the cell suspension, cells were lysed on ice with a Branson 450 Digital Sonifier (Branson Ultrasonics, Brookfield, CT) with 40% amplitude, for 5 min with 50% duty cycles. After 5 min incubation on ice, the sonification procedure was repeated. We the centrifuged the suspension 4 °C in a Beckman Coulter rotor Ti 45 at 30,000 rpm for 1 h, collected the supernatant and precipitated VP4 by adding 0.244 g ammonium sulfate per ml of protein solution. The resulting suspension was stirred in a glass flask overnight at 4 °C and 200 rpm and then spun at 4 °C in Beckman Coulter rotor Ti45 for 30 min at 30,000 rpm. We discarded the supernatant, resuspended the pellet in 60 ml of TE buffer (20 mM Tris, 1 mM EDTA, pH 8.0) with 1 mM PMSF, and transferred the solution to a Dounce homogenizer for homogenization with 15 strokes. The suspension was then spun at 4 °C for 30 min in a Beckman Coulter rotor Ti 45 at 30,000 rpm. Supernatant was collected and diluted with TE buffer with 1 mM PMSF to a final volume of 900 ml and loaded onto a 5 ml Q Sepharose column (Cytiva) using a peristaltic pump at a flow rate of 5 ml/min. Using an HPLC purification system, the column was washed with two CV of T10NE buffer (20 mM Tris, 10 mM NaCl, 1 mM EDTA, pH 8.0), and bound protein was eluted with 20 CV using a linear gradient from T10NE to T150NE (20 mM Tris, 150 mM NaCl, 1 mM EDTA, pH 8.0) at a flow rate of 1 ml/min. We pooled VP4-containing protein fractions, concentrated them to 500 μl using a 10 kDa MWCO centricon (Millipore Sigma), and exchanged the buffer to HNE with 0.1 mM PMSF using a G25 PD10 column. Protein was distributed to 420 μg aliquots and frozen in liquid nitrogen for storage at - 80 °C.

#### Liposome preparation

2 mg in total lipids of a mixture containing cholesterol, phosphocholine egg extract (egg PC), sphingomyelin (SM), 1,2-dioleyl-sn-glycero-3-phopshoethenolamine (DOPE), 1-palmitoyl-2-oleyo-sn-glycerol-3-phosphoetanolamine (POPE), 1,2-dioleyl-sn-glycero-3-phospho-L-serine (DOPS) and ganglioside GD1a (GD1a) in molar ratio of 50:12.5:10:8.5:8.5:8:2.5, dissolved in approximately 9:1 chloroform:methanol, were dried in a round bottom glass tube under argon flow. Remaining solvent was completely removed by incubating the lipids under high vacuum overnight. The lipids were resuspended in 200 μl of TNC buffer by vortexing for two minutes, in a volume corresponding to 10 mg/ml. We then freeze-thawed the lipid suspension twice using liquid nitrogen and formed liposomes by extrusion though a 0.2 μm filter using 41 passages. We evaluated the size of liposomes by dynamic light scattering using a 1:100 dilution of the liposome stock.

#### Recoating of DLPs with recombinant VP7 and VP4

We distributed 45 μg of DLPs in HNE equally among five 1.5 ml conical tubes (2.25 μl per tube). We first added 1 M sodium acetate, pH 5.2 to a final concentration of 100 mM and then added 65 μl VP4 (stored at 1.3 mg/ml) to a final concentration of 0.9 mg/ml in the final reaction volume, resulting in a 33-fold excess of VP4 monomer over a total of 180 sites on DLPs. A 0.1 mg/ml aprotinin solution was added to the samples to a final concentration of 0.2 μg/ml followed by incubation at 37 °C for 1 h. Required amounts of VP7 (9 μl stored at 1.0 mg/ml in HNE) to achieve a 2.3-fold excess of VP7 monomer over a total of 780 sites on DLPs were premixed with 0.1 volumes of TC buffer (20 mM Tris, 10 mM CaCl_2_, pH 8.0) and 0.1 volumes of 1 M sodium acetate pH 5.2 for 15 min before adding them to the DLP-VP4 mixture. Samples were incubated for 1 h at room temperature and then quenched by the addition of 0.1 volumes of 1 M Tris pH 8.0. Recoated particles from the five tubes were combined, and TNC was added to a final volume of 2.5 ml. rcTLPs were separated from excess VP4 and VP7 by ultracentrifugation at 4 °C in a Beckman Coulter rotor TLS 55 at 50,000 rpm for 1 h. We removed 2.0 ml of the supernatant, returned the volume to 2.5 ml with TNC, and pelleted again. Supernatant was carefully removed so that 100 to 200 μl remained. The rcTLP pellets were resuspended in the remaining buffer, sodium azide added to a final concentration of 0.01%, and the rcTLPs stored at 4 °C.

### Ca^2+^ permeabilization experiment and TIRF microscopy

#### Labelling of TLPs

TLPs were diluted to 1.0 mg/ml in a total volume of 30 μl using HNC (20 mM HEPES pH 8.0, 100 mM NaCl, 1 mM CaCl_2_) and 3.33 μl of 1 M NaHCO_3_ pH 8.3 added. This solution was mixed with 0.5 μl of 0.0076 mg/ml Atto 565 NHS ester. The reaction proceeded at room temperature for 1 h before quenching with 3 μl of 1 M Tris pH 8.0. The labeled TLPs were then buffer exchanged into a solution containing 20 mM Tris pH 8.0, 100 mM NaCl, and 1 mM CaCl_2_ using a Zeba Spin Desalting Column (Thermo Scientific).

#### Liposome preparation

A chloroform solution containing 2 mg total lipid mixture of cholesterol, phosphocholine egg extract (egg PC), sphingomyelin (SM), 1,2-dioleyl-sn-glycero-3-phopshoethenolamine (DOPE), biotinylated-DOPE, Cy5-DOPE, 1-palmitoyl-2-oleyo-sn-glycerol-3-phosphoetanolamine (POPE), 1,2-dioleyl-sn-glycero-3-phospho-L-serine (DOPS) and ganglioside GD1a (GD1a) in molar ratio of (40:22.5:10:7.25:0.5:0.25:8.5:8:2.5) was dried in a round bottom glass tube under argon flow. Remaining solvent was removed by incubating the lipids under high vacuum overnight. We then resuspended the lipids in 200 μl of TNE (20 mM Tris pH 8.0, 100 mM NaCl, 0.07 mM EGTA) containing 400 μM Fluo-4 dye by vortexing for two minutes, forming a 10 mg/ml lipid suspension, which we subjected to two freeze-thaw cycles with liquid nitrogen. By including 0.07 mM EGTA and then exchanging the liposomes into a buffer containing 1 mM CaCl_2_ before incubating them with TLPs (see below), we ensured that there was no free Ca^2+^ inside the liposomes. Liposomes were formed by extrusion with 41 passages through a 200 nm pore filter. The size of the liposomes was evaluated with dynamic light scattering using a 1:100 dilution of the liposome stock. The liposome suspension was loaded onto a G25 PD10 column equilibrated with TNC to separate liposomes from non-incorporated fluorophore and to exchange the buffer by eluting with 2 ml of TNC. We collected 500 μl fractions and used peak fractions containing liposomes for further experiments.

### Liposome Ca^2+^ permeabilization assay

4 μl of liposomes were added to 12 μg virus particles in a final volume of 18 μl and incubated for 30 min at 37 °C in the presence of 3.5 μg/ml trypsin. Trypsin was inactivated by adding aprotinin to a final concentration of 1 mg/ml. All samples were incubated for an additional 1.5 h at 37 °C. For controls, same amount of liposomes was incubated for the same time and temperature in the absence of virus (negative control) or in the presence of 20 μM ionomycin (positive control).

### TIRF measurements

25 mm circular glass coverslips were cleaned and coated with a 10% (w/v) solution of biotinylated-polyethylene glycol (PEG):PEG in a ratio 1:99 (Laysan Bio, cat. no. mPEG-SCM-5000) (Kyoung et al., 2013) and pre-incubated with 0.5 mg/ml neutrAvidin (Thermo Scientific, cat. no. LF144746) for 15 min at room temperature (RT) before use. Liposome solutions were diluted 500 times in TNC in order to avoid stochastic co-localizations of virus and liposomes, incubated over the coverslip for 10 min at RT, and unbound liposomes were removed before imaging by a single wash with 100 μl TNC. Images of different fields of view were acquired under TIRF illumination using an Olympus IX70 microscope equipped with a Hamamatsu ImageEM camera and excitation from 488, 561 and 640 nm lasers (Coherent, Inc., Salem, NH) operated at 100, 150 and 70 mW, respectively. For each field of view, we recorded three images with 50% power from each laser and 30 ms exposure at 488 nm (Fluo-4 dye, Ca^2+^ sensor signal), 50 ms exposure at 561 nm (Atto 565 NHS ester dye, virus signal), 20 ms exposure at 640 nm (Cy5-DOPE dye, liposome signal). The microscope was controlled by the SlideBook acquisition program (Intelligent Imaging Innovations, Denver, CO), and image processing was carried out using Fiji software (Schindelin et al., 2012).

### Image analysis

We used custom-made MATLAB (MathWorks) scripts for automatic detection of fluorescent liposomes in the 640 nm channel (Cy5-DOPE dye, liposome signal) images by fitting a 2D Gaussian function to diffraction limited spots for which the intensity was 1.5 above background. We extracted the XY position of each liposome in all images and calculated the integrated fluorescence intensity on a 4x4 pixel area around the XY position for the 3 channels. We calculated the background by measuring the integrated fluorescence intensity on a 6x6 pixel region around the XY position, which was then was subtracted from the integrated intensity on the 4x4 area after area correction for the 3 different channels. For the samples that were incubated with TLPs, the liposomes that colocalized with virus were selected by applying a threshold on the integrated fluorescence intensity in the 561 channel (Atto 565 NHS ester dye, virus signal) at the same XY position; the threshold was calculated with images, taken at the same time, that contained only virus. We measured the integrated intensity of diffraction limited spots by fitting 2D Gaussian functions. We used a t-test to identify valid spots for which the signal was statistically higher than the local background.

## Cryo-EM specimen preparation and data collection

### Incubation of rcTLPs with liposomes

To rcTLP stock solutions (0.7–1.5 mg/ml), we added liposome stock solutions (10 mg/ml, average diameter 200 nm) to a final concentration of 1.5 mg/ml, followed by the addition of trypsin to a final concentration of 5 μg/ml and incubation for 30 min at 37 °C. Trypsin was then inactivated with aprotinin at a final concentration of 1 mg/ml. Samples were incubated at 37 °C for an additional 30 min.

### Cryo-EM sample preparation

C-flat holey copper carbon grids (CF-2/1-2C, Electron Microscopy Sciences) were glow discharged for 30 s at 10 mA. Using a CP3 cryo-plunger, 4 μl of virus-liposome samples were applied to the grids, blotted for 4 s with at a humidity ranging from 88% to 92% and plunged into liquid ethane having a temperature of -273 °C. Grids were stored in liquid nitrogen.

### Cryo-EM data collection

We recorded movies with a Titan Krios G3i microscope (Thermo Fisher Scientific, Waltham, MA), operated at 300 kV and equipped with a BioQuantum energy filter and a K3 direct electron detector camera (Gatan, Inc.). Dose-fractionated movies with 50 frames were recorded in counting mode with 0.05 s per frame for a total of 2.5 s with SerialEM v3.7 (Mastronarde, 2005). The magnification was 60,606, resulting in a pixel size of 0.825 Å per pixel. For some grids, we recorded a total of 27 movies for each stage position (nanoprobe mode with an illuminated area of ∼1 μm, 3 exposures per hole from a total of 9 holes).

## Cryo-EM data processing

### Preprocessing and initial particle stack preparation

We used MotionCor2 (Zheng et al., 2017) with 5×5 patch alignment to obtain summed micrographs from the frames of 33,563 movies that we had recorded from four sample preparations. With CTFFIND4 (Rohou and Grigorieff, 2015), we estimated contrast transfer function (CTF) parameters from the summed micrographs. We then picked 77,139 rotavirus particles from the micrographs template matching in cisTEM (Grant et al., 2018; Lucas et al., 2021), with the routines match_template (version 1.00), refine_template (version 1.00), and make_template_result (version 1.00). We carried out template matching at 5 Å resolution with angular search steps of 1.7° and a previously determined reconstruction as reference structure (Herrmann et al., 2021), resampled to the correct pixel size. The symmetry for the search was I_2_ and the radius of the mask 403 Å. We determined CTF parameter values at particle positions with Gctf (Zhang, 2016) and used relion_preprocess (Scheres, 2012) for particle extraction and normalization. The box size was 1536×1536 pixels and the radius for background area designation 666 pixels. After 2D classification with relion_refine (Scheres, 2012), we kept 75,846 particle images for analysis.

### Icosahedral reconstruction

An initial reconstruction calculated with the alignment parameters obtained from template matching had a nominal resolution of 3.89 Å as determined by Fourier shell correlation (FSC) with a cutoff value of 0.143 and after masking the half maps with a spherical shell mask that had inner and outer radii of 222 and 403 Å, respectively (Figure S4). We then improved particle alignment, particle images, and reconstruction parameters by iterating routines in cisTEM (refine3d version 1.01, reconstruct3d version 1.02) (Grant et al., 2018) with routines in RELION (relion_motion_refine, and relion_ctf_refine) (Scheres, 2014). The resolution for alignment in the last iteration was 3.4 Å. In relion_ctf_refine, we used per-particle defocus fitting, beam-tilt refinement, symmetric aberration estimation, and fitting of anisotropic magnification distortion for each of the particle stack’s 63 optics groups (--fit_defocus –kmin_defocus 30 –fit_mode fpmfp –fit_beamtilt –kmin_tilt 30 –fit_aniso – odd_aberr_max_n 3 –fit_aberr). Reconstuctions were calculated with relion_reconstruct with the option –fom_weighting, where we assigned a per particle figure of merit, *FOM*, based on the cisTEM alignment scores, where *S_i_* is the score of particle *I*, and *S_max_* and *S_min_* are the highest and lowest scores, respectively:

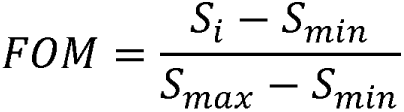

After two iterations, the nominal resolution of the final icosahedral reconstruction of full viral particles converged to 2.43 Å as estimated with relion_postprocess (Table S1 and Figure S4).

### Local analysis of spike positions

For local analysis of individual spike positions, we used protocols similar to those previously described (Herrmann et al., 2021; Jenni et al., 2022; Jenni et al., 2019), with some modifications (Figure S5). Generally the workflow for preparing subparticle stacks for local analysis involved symmetry expansion of the RELION particle star file, followed by signal subtraction with relion_project using a suitable mask to retain density, and sub-particle extraction from the signal-subtracted images with custom Python scripts and IMOD routines (Mastronarde and Held, 2017). Depending on the analysis, different subparticle stacks were prepared.

For initial classification of spike positions -- that is, determining for each of the 60 positions on the virion whether the VP5*/VP8* penetration protein was in an upright, intermediate or reversed conformation, or whether it was entirely absent -- we made a 384×384 pixel subparticle stack 1 in which we had signal-subtracted all density except within the volume of a single VP7 trimer plus the volume potentially occupied by VP5*/VP8* in upright and reversed conformations (Figure S5A–D). We carried out classifications with refine3d and reconstruct3d from cisTEM (Grant et al., 2018), keeping the subparticle alignment angles and shifts constant during all iterations. We also turned off per-particle weighting in the reconstruction step by setting the particle weighting factor input parameter of reconstruct3d to 0.0 and by resetting the LogP, SIGMA, and SCORE values in par files to constant values of -2000, 4.1, and 20.1, respectively, and running the reconstruction step again before proceeding to the next iteration. Initial classes were seeded randomly. In a first classification (classification #1), we asked for 16 classes (Table S2). The resolution limit for classification was 5 Å and the radius of the spherical mask applied to the 2D images was 140 Å. 3D references were masked around the single VP7 trimer and VP5*/VP8* volume. Inspection of the resulting classes from the first classification showed that particles partitioned not only based on spike occupancy and conformation, but also based on radial shifts (up to approximately 5 Å) with respect to the virion center (Movie S1). As evident from Figure S6, the main cause for the observed radial shifts appeared to be anisotropic magnification distortion, because the class assignment of subparticles from the two classes with the largest relative shift observed (classes 4 and 13) showed a systematic bias depending on the extraction angle with respect to the virion center. We corrected for anisotropic magnification when we calculated icosahedral reconstructions of full viral particles, but no correction was implemented during subparticle extraction. Other imperfections probably also contributed to the observed shifts; for instance, if data had been collected slightly away from the eucentric height of the microscope. We therefore further classified each of the 16 classes in a second classification (classification #2), aligned all 32 classes by rigid-body fitting a model of the VP7 trimer and updating the alignment parameters in the RELION star file based on the superpositions of the fitted models, and made a final assignment of each subparticle as upright, intermediate, reversed, or unoccupied (Figure S5E–F, Figure S7, and Table S2).

### Local analysis of reversed VP5*/VP8* spikes bound to liposomes

Because we observed minor density artifacts in reconstructions calculated from subparticle stack 1 (see above) when contoured at very low level (presumably caused by a bug or array overflow in relion_project when the volume for signal subtraction had a large size of 1536×1536×1536 pixels), we prepared subparticle stack 2 with slightly down sampled data of 1024×1024×1024 pixels for the full virus particle box, corresponding to a pixel size of 1.2375 Å. For this stack, we signal-subtracted all density within a tight-fitting spherical mask except within the volume occupied by VP5*/VP8* at the reference icosahedral position (Figure S5C–D). Classifications #3 and #4 were carried out starting with 1,510,279 particles in reversed conformation selected from the signal-subtracted subparticle stack 2 based on the results from classification #1 and #2 (Figure S5G). 2D and 3D masking was with a radius of 124 Å, and the resolution for classification was 5 and 8 Å, respectively. Subparticles from classes 1 and 3 of classification #3 that showed liposome density were merged and subjected to classification #4 (Figure S8 and Table S3). Final reconstructions of liposome-bound classes were computed from the original (no signal subtraction) subparticle stack 2 (Figure S5H).

### Mapping of subparticles in the original micrographs

When we extracted subparticles, we assigned unique viral particle and subparticle identifiers together with the *x* and y coordinates of the extraction position of each subparticle with respect to the original extraction position of the corresponding virus in the micrograph. We could thus map subparticles from liposome-bound classes onto the original micrographs (Figure 3D).

## Supporting information

Supplemental animation

## Figure preparation

We prepared the figures using PyMOL (The PyMOL Molecular Graphics System, Version 2.3 Schrödinger, LLC), ChimeraX (Pettersen et al., 2021), BioRender, matplotlib (Hunter, 2007) and ImageMagick (ImageMagick Studio LLC, 2023, available at: https://imagemagick.org).

## Data availability

The cryo-EM maps are deposited in the Electron Microscopy Data Bank with accession identifiers EMD-42343 (class 5) and EMD-42344 (class 6). Atomic coordinates are deposited in the Protein Data Bank with accession identifiers PDB-ID 8UK2 (class 5) and PDB-ID 8UK3 (class 6).

## ACKNOWLEDGMENTS

We thank Gustavo Scanavachi (PCMM, Boston Children’s Hospital) for essential help with fluorescence microscopy and image analysis, Tom Kirchhausen (PCMM, Boston Children’s Hospital and Dept. of Cell Biology, Harvard Medical School) for access to up-to-date microscopy and image-analysis facilities, and the staff at the Harvard Medical School Center for Cryo-Electron Microscopy in Structural Biology, for expert and effective assistance. Partial support for the Center came from the Nancy Lurie Marks Foundation. A grant from NIH, R01 CA13202, supported the research.

## AUTHOR CONTRIBUTIONS

MDS and TH prepared reagents, recorded and curated data, analyzed data, prepared figures, and participated in drafting and editing the manuscript; SJ curated and analyzed data, prepared figures, supervised research, and participated in drafting and editing the MS; SCH supervised research and participated in drafting and editing the manuscript.

## FIGURE CAPTIONS

**Figure S1. Single-particle fluorescence imaging.** (A) Fluorescence intensity for each liposome in three independent experiments (Experiment 1, 2 and 3). Fluorescence intensity corresponding to the control sample (left), the ionophore sample (center) or TLP sample (right) are shown for each experiment, respectively. Top graph (cyan) shows the signal in the 640nm channel (Cy5-DOPE dye, liposome signal); center graph (green) shows the signal in the 488 nm channel (Fluo-4 dye, Ca^2+^ sensor signal); and bottom graph (magenta) shows the signal in 561 nm channel (Atto 565 dye, TLP signal). (B) Normalized frequency of the liposome signal for the control sample (white bars), ionophore sample (light grey bars) and TLP sample (all liposomes, dark grey bars) in each experiment.

**Figure S2. Ca^2+^-signal analysis from the single-particle fluorescence imaging.** (A) Ca^2+^ signal as a function of liposome signal for the control sample (white dots) and for ionophore sample (light grey dots) in the experiment 1 (left), experiment 2 (center) and experiment 3 (right), respectively. (B) Ca^2+^ signal as a function of the binned liposome signal for the control sample (white bars) and the ionophore sample (light grey bars) in experiment 1, 2 and 3, respectively. Error bars represent the standard deviation calculated from individual measurements within each bin as shown in the scatter plot in panel A. (C) Ca^2+^ signal as a function of liposome signal for TLP control (small random selection of liposomes that did not colocalize with virus in the TLP sample, white dots) and for TLP virus (liposomes colocalizing with virus only, dark grey dots) samples in the experiment 1 (left), experiment 2 (center) and experiment 3 (right), respectively. (D) Ca^2+^ signal as a function of the binned liposome signal for the TLP control sample (liposomes that do not colocalize with virus in the TLP sample, white bars) and the TLP virus sample (liposomes colocalizing with virus only, dark grey bars) in experiments 1, 2 and 3, respectively. Error bars represent the standard deviation calculated from individual measurements within each bin as shown in the scatter plot in panel C. Statistical analysis in B and D used a t-test between samples. n.s = p>0.05; * = p<0.05; ** = p<0.01; *** = p<0.001.

**Figure S3. Cryo-EM sample preparation of liposome-bound rotaviruses.** Schematic illustration of the interaction between rotavirus and a liposome during specimen preparation for cryo-EM analysis.

**Figure S4. Icosahedral reconstruction of RRV particles.** (A) Fourier shell correlations (FSCs) between half maps calculated from icosahedral reconstructions after applying a spherical shell mask encompassing the three protein layers of the TLP (VP2, VP6, and VP7). Curves are shown for the data processing steps as described in Methods. The final nominal resolution was 2.43 Å. (B) Local resolution analysis of the icosahedral reconstruction. A segment of the “triple-layer” particle is shown. (C) Density maps of VP2, VP6 and VP7 regions.

**Figure S5. Cryo-EM analysis processing scheme of TLPs with liposomes.** (A) Alignment of full viral particles with icosahedral symmetry imposed. A map of the final icosahedral reconstruction is shown where the particle is partially cut. VP1 (RNA-dependent RNA polymerase) and RNAs are colored gray; VP2, cyan and blue; VP6, green; VP7, yellow. See also Figure S4. (B) Icosahedral symmetry expansion (60-fold) based on the full particle alignment. (C) Signal subtraction. The masks used to the define the subtraction volume, excluding the region of interest at a single protomer position, are shown. Signal-subtracted stack 1 was used to prepare subparticles for classifications #1 and #2 (spike conformation and occupancy). Signal-subtracted stack 2 was used to prepare subparticles for classifications #3 and #4 (liposome interaction of reversed spikes). (D) Subparticle extraction from the signal-subtracted and original particles stacks. (E) Classifications #1 and #2, see Figure S7. (F) Alignment of classes to correct for observed shifts in reconstructions of subparticle classes caused primarily by anisotropic magnification distortion, see Figure S6. Subparticle reconstructions are shown for the three observed spike conformations (upright, intermediate, and reversed) and empty positions. Density maps were low pass filtered at 5 Å resolution and partially cut. VP2, cyan and blue; VP6, green; VP7, yellow; VP5*, red, orange, and salmon. (G) Classifications #3 and #4, see Figure S8. (H) Final reconstructions of liposome-bound classes calculated from subparticle stack 2, which was obtained from the original (non-signal-subtracted) images.

**Figure S6. Observed subparticle shifts caused by anisotropic magnification distortion.** (A) Subparticles were classified without alignment. Relative shifts between classes were determined by rigid-body fitting a V7 trimer model and then aligning the classes by updating the subparticle alignment parameters based on the fitted models. (B) Ribbon representation of the VP7 trimer models after fitting it into the densities of the 32 classes. The largest shift we observed was 5.7 Å between classes 4 and 13. (C) Each subparticle is associated with an extraction radius and angle with respect to the center of the full virus particle projection in the original micrograph. (D) Histograms of metadata values (extraction radius, extraction angle, defocus, optics group number, micrograph number) for all subparticles from the two classes 4 and 13, which showed the largest relative shift in their reconstructions. A strong bias in the extraction angle suggests that the observed shifts are predominately caused by anisotropic magnification distortion, which was not accounted for during subparticle extraction. (D) Histograms of metadata values (extraction radius, extraction angle, defocus, optics group number, micrograph number) for dataset 2 subparticles from the two classes 4 and 13.

**Figure S7. Classification #1 and #2.** For display, density maps were calculated from subparticle stack 1, which was obtained from the original (non-signal-subtracted) images, low pass filtered at 5 Å resolution and partially cut. VP2, cyan and blue; VP6, green; VP7, yellow; VP5*, red, orange, and salmon. (A) Classification #1 partitioned the particles into 16 classes, each of which was further subclassified into two classes in classification #2 (indicated by dashed lines). (B) Reconstructions of three observed spike conformations (upright, intermediate, and reversed) and empty positions after merging corresponding classes (Table S2) and correcting of observed subparticle shifts (Figure S6).

**Figure S8. Classification #3 and #4.** Classification of liposome-bound spike positions from subparticles with reversed spike conformations. (A) In classification #3, we initially requested four classes. Density maps were calculated from signal-subtracted subparticle stack 2, low pass filtered at 8 Å resolution and displayed in gray and at very low contour level to visualize membrane density. (B) In classification #4, we further subclassified classes 1 and 3 from classification #3. Density maps of the six classes were calculated from subparticle stack 2, which was obtained from the original (non-signal-subtracted) images, and low pass filtered at 8 Å. VP6, green; VP7, yellow; VP5*, red, orange, and salmon; membrane, gray.

**Figure S9. Local resolution analysis of liposome-bound VP5*.** (A) Full view of the class 5 and class 6 reconstructions colored according to local resolution. (B) Corresponding slab views.

## REFERENCES

Abdelhakim, A.H., Salgado, E.N., Fu, X., Pasham, M., Nicastro, D., Kirchhausen, T., and Harrison, S.C. (2014). Structural correlates of rotavirus cell entry. PLoS Pathog 10, e1004355.

Agosto, M.A., Ivanovic, T., and Nibert, M.L. (2006). Mammalian reovirus, a nonfusogenic nonenveloped virus, forms size-selective pores in a model membrane. Proc. Natl. Acad. Sci. U. S. A. 103, 16496–16501.

Altenburg, B.C., Graham, D.Y., and Estes, M.K. (1980). Ultrastructural study of rotavirus replication in cultured cells. J. Gen. Virol. 46, 75–85.

Aoki, S.T., Settembre, E.C., Trask, S.D., Greenberg, H.B., Harrison, S.C., and Dormitzer, P.R. (2009). Structure of rotavirus outer-layer protein VP7 bound with a neutralizing Fab. Science 324, 1444–1447.

Bergmann, C.C., Maass, D., Poruchynsky, M.S., Atkinson, P.H., and Bellamy, A.R. (1989). Topology of the non-structural rotavirus receptor glycoprotein NS28 in the rough endoplasmic reticulum. EMBO J. 8, 1695–1703.

Crawford, S., Ding, S., Greenberg, H.B., and Estes, M.K. (2023). Rotaviruses. In Fields Virology, 7th ed., P.M. Howley, and D.M. Knipe, eds. (Philadelphia, PA: Wolters Kluyver), pp. 362-413.

Delorme, C., Brussow, H., Sidoti, J., Roche, N., Karlsson, K.A., Neeser, J.R., and Teneberg, S. (2001). Glycosphingolipid binding specificities of rotavirus: identification of a sialic acid-binding epitope. J. Virol. 75, 2276–2287.

Ding, K., Celma, C.C., Zhang, X., Chang, T., Shen, W., Atanasov, I., Roy, P., and Zhou, Z.H. (2019). In situ structures of rotavirus polymerase in action and mechanism of mRNA transcription and release. Nat. Commun. 10, 2216.

Dormitzer, P.R., Sun, Z.Y., Blixt, O., Paulson, J.C., Wagner, G., and Harrison, S.C. (2002). Specificity and affinity of sialic acid binding by the rhesus rotavirus VP8* core. J. Virol. 76, 10512–10517.

Grant, T., Rohou, A., and Grigorieff, N. (2018). cisTEM, user-friendly software for single-particle image processing. Elife 7, e35383.

Greenberg, H.B., Valdesuso, J., van Wyke, K., Midthun, K., Walsh, M., McAuliffe, V., Wyatt, R.G., Kalica, A.R., Flores, J., and Hoshino, Y. (1983). Production and preliminary characterization of monoclonal antibodies directed at two surface proteins of rhesus rotavirus. J. Virol. 47, 267–275.

Helfrich, W. (1973). Elastic properties of lipid bilayers: theory and possible experiments. Z Naturforsch C 28, 693–703.

Herrmann, T., Torres, R., Salgado, E.N., Berciu, C., Stoddard, D., Nicastro, D., Jenni, S., and Harrison, S.C. (2021). Functional refolding of the penetration protein on a non-enveloped virus. Nature 590, 666–670.

Hu, L., Crawford, S.E., Czako, R., Cortes-Penfield, N.W., Smith, D.F., Le Pendu, J., Estes, M.K., and Prasad, B.V. (2012). Cell attachment protein VP8* of a human rotavirus specifically interacts with A-type histo-blood group antigen. Nature 485, 256–259.

Hunter, J.D. (2007). Matplotlib: A 2D graphics environment. Comput. Sci. Eng. 9, 90–95.

Jenni, S., Li, Z., Wang, Y., Bessey, T., Salgado, E.N., Schmidt, A.G., Greenberg, H.B., Jiang, B., and Harrison, S.C. (2022). Rotavirus VP4 Epitope of a Broadly Neutralizing Human Antibody Defined by Its Structure Bound with an Attenuated-Strain Virion. J. Virol. 96, e0062722.

Jenni, S., Salgado, E.N., Herrmann, T., Li, Z., Grant, T., Grigorieff, N., Trapani, S., Estrozi, L.F., and Harrison, S.C. (2019). In situ Structure of Rotavirus VP1 RNA-Dependent RNA Polymerase. J. Mol. Biol. 431, 3124–3138.

Kim, I.S., Trask, S.D., Babyonyshev, M., Dormitzer, P.R., and Harrison, S.C. (2010). Effect of mutations in VP5 hydrophobic loops on rotavirus cell entry. J. Virol. 84, 6200–6207.

Kumar, D., Yu, X., Crawford, S.E., Moreno, R., Jakana, J., Sankaran, B., Anish, R., Kaundal, S., Hu, L., Estes, M.K., et al. (2020). 2.7 A cryo-EM structure of rotavirus core protein VP3, a unique capping machine with a helicase activity. Sci Adv 6, eaay6410.

Kyoung, M., Zhang, Y., Diao, J., Chu, S., and Brunger, A.T. (2013). Studying calcium-triggered vesicle fusion in a single vesicle-vesicle content and lipid-mixing system. Nat Protoc 8, 1–16.

Lucas, B.A., Himes, B.A., Xue, L., Grant, T., Mahamid, J., and Grigorieff, N. (2021). Locating macromolecular assemblies in cells by 2D template matching with cisTEM. Elife 10.

Martinez, M.A., Lopez, S., Arias, C.F., and Isa, P. (2013). Gangliosides have a functional role during rotavirus cell entry. J. Virol. 87, 1115–1122.

Mastronarde, D.N. (2005). Automated electron microscope tomography using robust prediction of specimen movements. J. Struct. Biol. 152, 36–51.

Mastronarde, D.N., and Held, S.R. (2017). Automated tilt series alignment and tomographic reconstruction in IMOD. J. Struct. Biol. 197, 102–113.

Ogden, K.M., Snyder, M.J., Dennis, A.F., and Patton, J.T. (2014). Predicted structure and domain organization of rotavirus capping enzyme and innate immune antagonist VP3. J. Virol. 88, 9072–9085.

Pettersen, E.F., Goddard, T.D., Huang, C.C., Meng, E.C., Couch, G.S., Croll, T.I., Morris, J.H., and Ferrin, T.E. (2021). UCSF ChimeraX: Structure visualization for researchers, educators, and developers. Protein Sci. 30, 70–82.

Poruchynsky, M.S., and Atkinson, P.H. (1991). Rotavirus protein rearrangements in purified membrane-enveloped intermediate particles. J. Virol. 65, 4720–4727.

Ramani, S., Cortes-Penfield, N.W., Hu, L., Crawford, S.E., Czako, R., Smith, D.F., Kang, G., Ramig, R.F., Le Pendu, J., Prasad, B.V., et al. (2013). The VP8* domain of neonatal rotavirus strain G10P[11] binds to type II precursor glycans. J. Virol. 87, 7255–7264.

Rohou, A., and Grigorieff, N. (2015). CTFFIND4: Fast and accurate defocus estimation from electron micrographs. J. Struct. Biol. 192, 216–221.

Salgado, E.N., Garcia Rodriguez, B., Narayanaswamy, N., Krishnan, Y., and Harrison, S.C. (2018). Visualization of Calcium Ion Loss from Rotavirus during Cell Entry. J. Virol. 92, e01327–01318.

Salgado, E.N., Upadhyayula, S., and Harrison, S.C. (2017). Single-particle detection of transcription following rotavirus entry. J. Virol., e00651–00617.

Scheres, S.H. (2012). RELION: implementation of a Bayesian approach to cryo-EM structure determination. J. Struct. Biol. 180, 519–530.

Scheres, S.H. (2014). Beam-induced motion correction for sub-megadalton cryo-EM particles. Elife 3, e03665.

Schindelin, J., Arganda-Carreras, I., Frise, E., Kaynig, V., Longair, M., Pietzsch, T., Preibisch, S., Rueden, C., Saalfeld, S., Schmid, B., et al. (2012). Fiji: an open-source platform for biological-image analysis. Nat. Methods 9, 676-682.

Settembre, E.C., Chen, J.Z., Dormitzer, P.R., Grigorieff, N., and Harrison, S.C. (2011). Atomic model of an infectious rotavirus particle. EMBO J. 30, 408–416.

Shah, P.N.M., Gilchrist, J.B., Forsberg, B.O., Burt, A., Howe, A., Mosalaganti, S., Wan, W., Radecke, J., Chaban, Y., Sutton, G., et al. (2023). Characterization of the rotavirus assembly pathway in situ using cryoelectron tomography. Cell Host Microbe 31, 604–615 e604.

Soler, C., Musalem, C., Lorono, M., and Espejo, R.T. (1982). Association of viral particles and viral proteins with membranes in SA11-infected cells. J. Virol. 44, 983–992.

Tihova, M., Dryden, K.A., Bellamy, A.R., Greenberg, H.B., and Yeager, M. (2001). Localization of membrane permeabilization and receptor binding sites on the VP4 hemagglutinin of rotavirus: implications for cell entry. J. Mol. Biol. 314, 985–992.

Trask, S.D., Kim, I.S., Harrison, S.C., and Dormitzer, P.R. (2010). A rotavirus spike protein conformational intermediate binds lipid bilayers. J. Virol. 84, 1764–1770.

Trask, S.D., McDonald, S.M., and Patton, J.T. (2012). Structural insights into the coupling of virion assembly and rotavirus replication. Nat. Rev. Microbiol. 10, 165–177.

Xia, X., Wu, W., Cui, Y., Roy, P., and Zhou, Z.H. (2021). Bluetongue virus capsid protein VP5 perforates membranes at low endosomal pH during viral entry. Nat Microbiol 6, 1424–1432.

Zhang, K. (2016). Gctf: Real-time CTF determination and correction. J. Struct. Biol. 193, 1–12.

Zhang, L., Agosto, M.A., Ivanovic, T., King, D.S., Nibert, M.L., and Harrison, S.C. (2009). Requirements for the formation of membrane pores by the reovirus myristoylated micro1N peptide. J. Virol. 83, 7004-7014.

Zhang, X., Patel, A., Celma, C.C., Yu, X., Roy, P., and Zhou, Z.H. (2016). Atomic model of a nonenveloped virus reveals pH sensors for a coordinated process of cell entry. Nat. Struct. Mol. Biol. 23, 74–80.

Zheng, S.Q., Palovcak, E., Armache, J.P., Verba, K.A., Cheng, Y., and Agard, D.A. (2017). MotionCor2: anisotropic correction of beam-induced motion for improved cryo-electron microscopy. Nat. Methods 14, 331–332.

