## Supplemental figures and tables for "The rotavirus VP5*/VP8* conformational transition permeabilizes membranes to Ca^2+^"

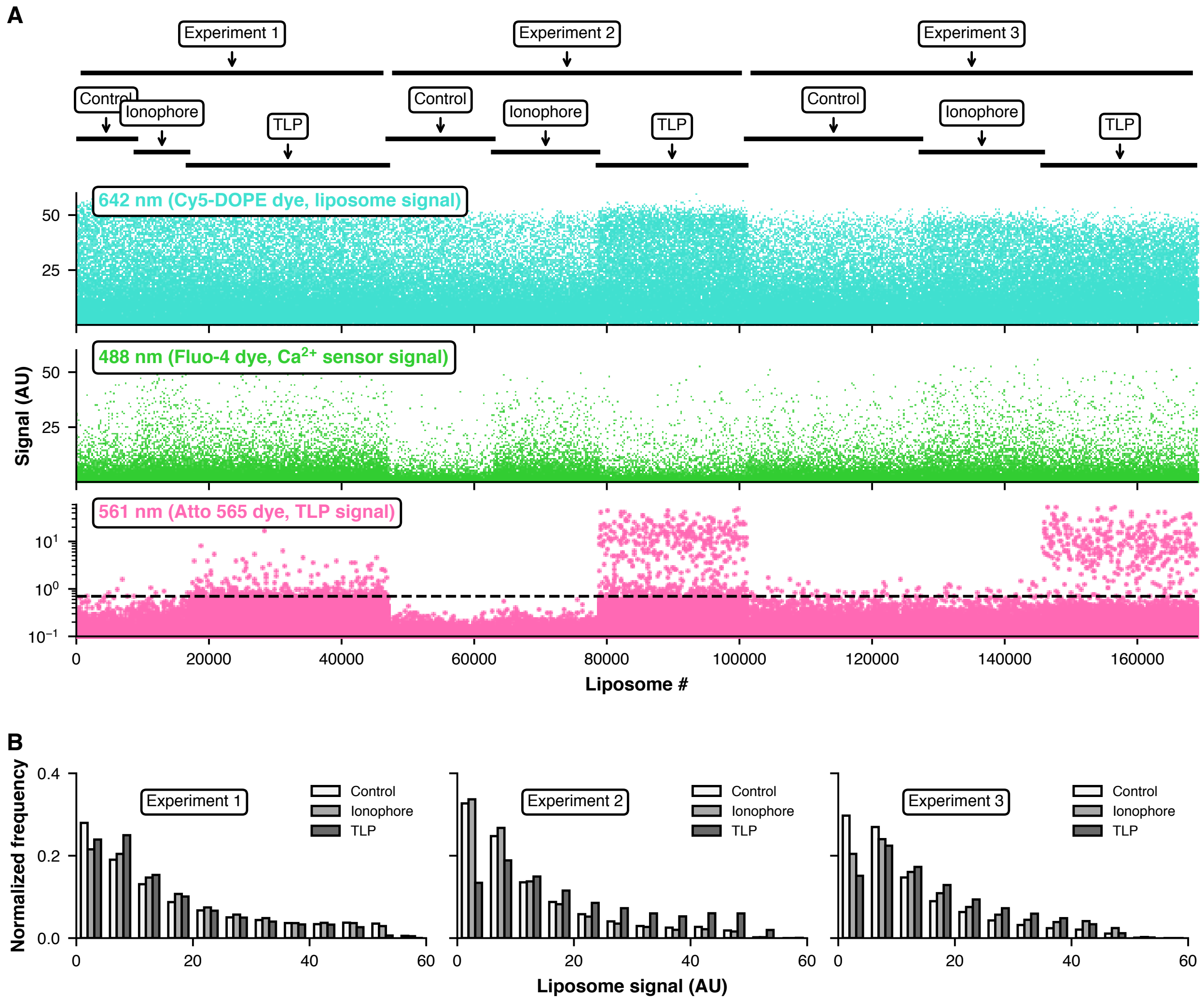


**Figure S1. Single-particle fluorescence imaging.**

(A) Fluorescence intensity for each liposome in three independent experiments (Experiment 1, 2 and 3). Fluorescence intensity corresponding to the control sample (left), the ionophore sample (center) or TLP sample (right) are shown for each experiment. Top graph (cyan) shows the signal in the 642 nm channel (Cy5-DOPE dye, liposome signal); center graph (green) shows the signal in the 488 nm channel (Fluo-4 dye, $Ca^{2+}$ sensor signal); and bottom graph (magenta) shows the signal in 561 nm channel (Atto 565 dye, TLP signal). (B) Normalized frequency of the liposome signal for the control sample (white bars), ionophore sample (light grey bars) and TLP sample (all liposomes, dark grey bars) in each experiment.

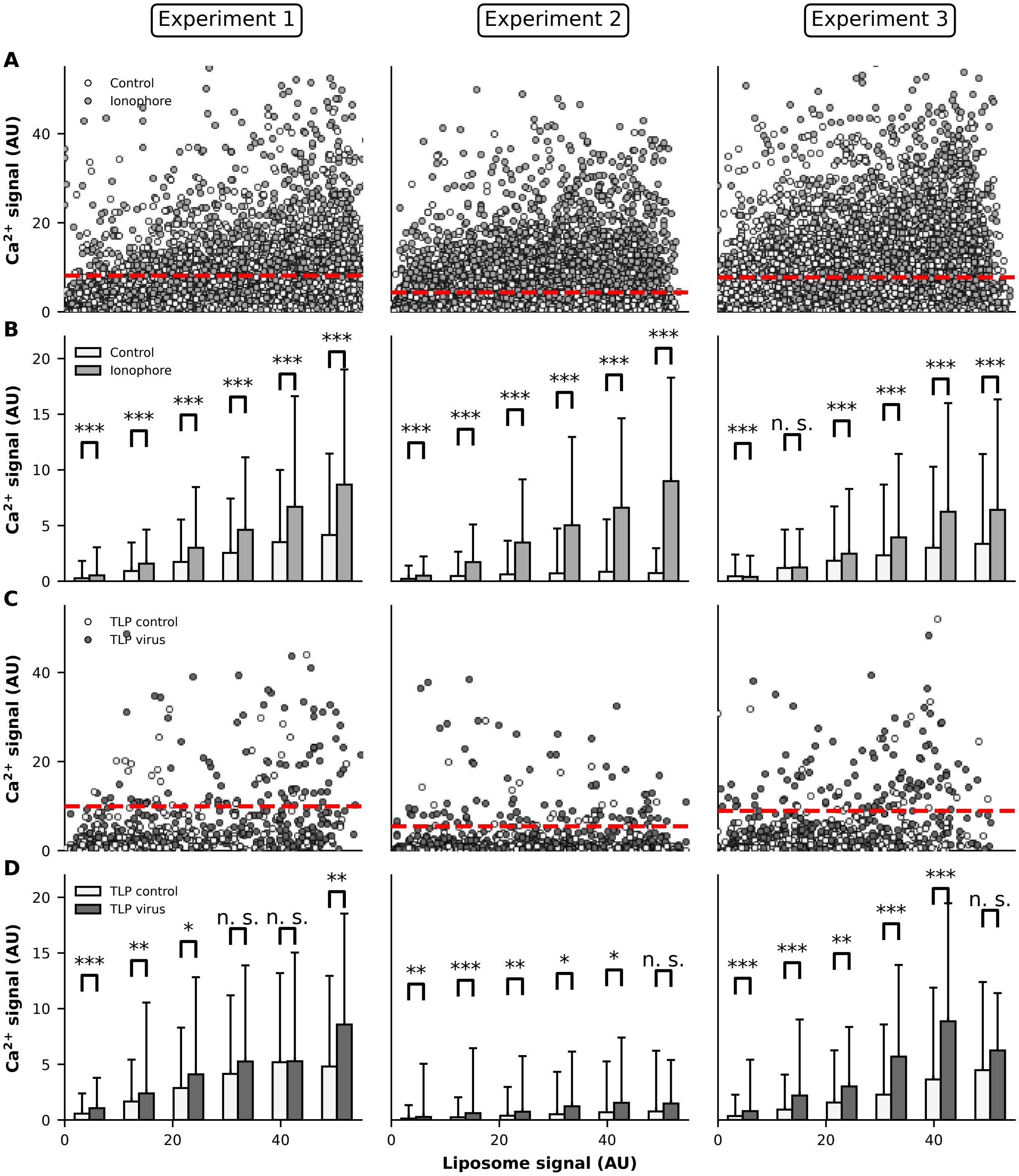


**Figure S2. $Ca^{2+}$-signal analysis from the single-particle fluorescence imaging.**

(A) $Ca^{2+}$ signal as a function of liposome signal for the control sample (white dots) and for ionophore sample (light grey dots) in the experiment 1 (left), experiment 2 (center) and experiment 3 (right), respectively. (B) $Ca^{2+}$ signal as a function of the binned liposome signal for the control sample (white bars) and the ionophore sample (light grey bars) in experiment 1, 2 and 3, respectively. Error bars represent the standard deviation calculated from individual measurements within each bin as shown in the scatter plot in panel A. (C) $Ca^{2+}$ signal as a function of liposome signal for TLP control (small random selection of liposomes that do not colocalize with virus in the TLP sample, white dots) and for TLP virus (liposomes colocalizing with virus only, dark grey dots) samples in the experiment 1 (left), experiment 2 (center) and experiment 3 (right), respectively. (D) $Ca^{2+}$ signal as a function of the binned liposome signal for the TLP control sample (liposomes that do not colocalize with virus in the TLP sample, white bars) and the TLP virus sample (liposomes colocalizing with virus only, dark grey bars) in experiment 1, 2 and 3, respectively. Error bars represent the standard deviation calculated from individual measurements within each bin as shown in the scatter plot in panel C. Statistical analysis was performed in B and D using a t-test between samples. n.s = $p>0.05$; * = $p<0.05$; ** = $p<0.01$; *** = $p<0.001$.

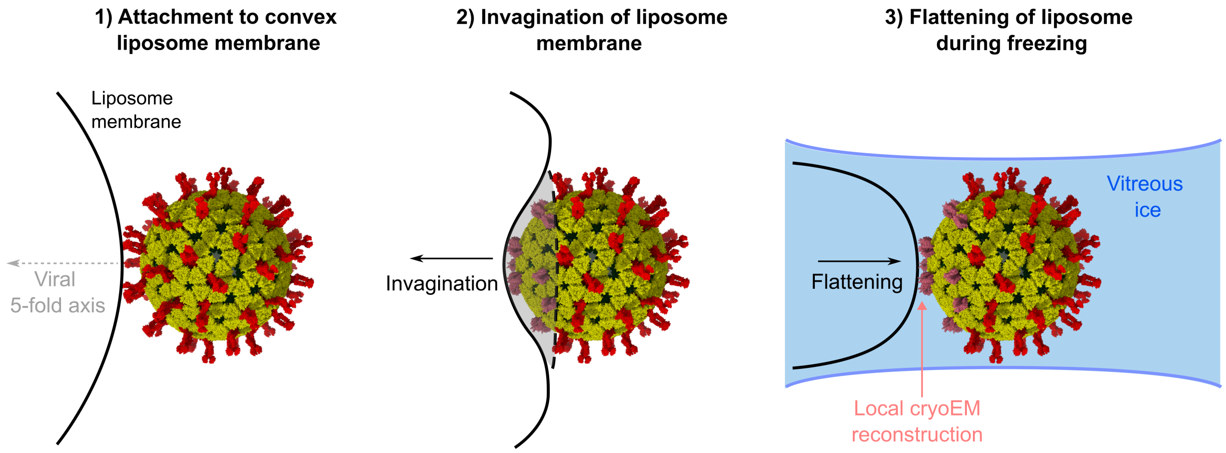


**Figure S3. Cryo-EM sample preparation of liposome-bound rotaviruses.**
Schematic illustration of the interaction between rotavirus and a liposome during specimen preparation for cryo-EM analysis.

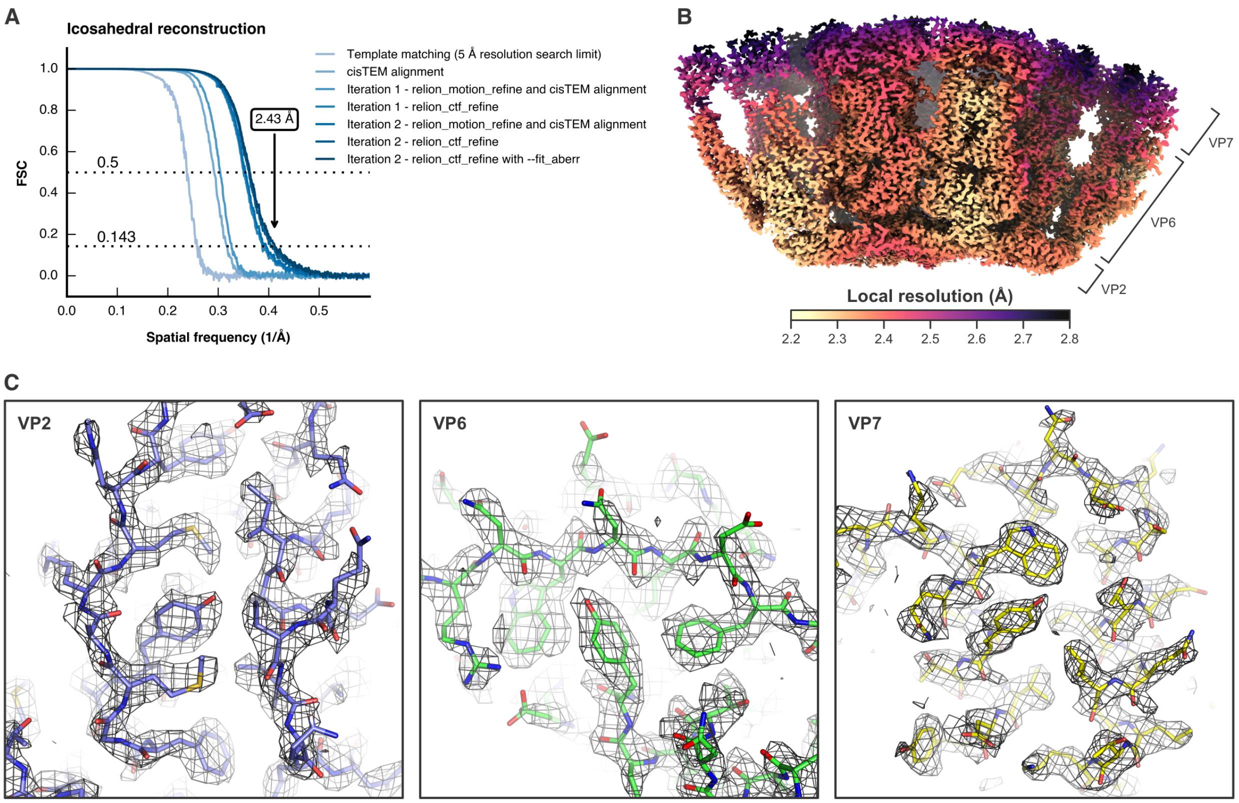


**Figure S4. Icosahedral reconstruction of RRV particles.**

(A) Fourier shell correlations (FSCs) between half maps calculated from icosahedral reconstructions after applying a spherical shell mask encompassing the three protein layers of the TLP (VP2, VP6, and VP7). Curves are shown for the data processing steps as described in Methods. The final nominal resolution was 2.43 Å. (B) Local resolution analysis of the icosahedral reconstruction. A segment of the "triple-layer" particle is shown. (C) Density maps of VP2, VP6 and VP7 regions.

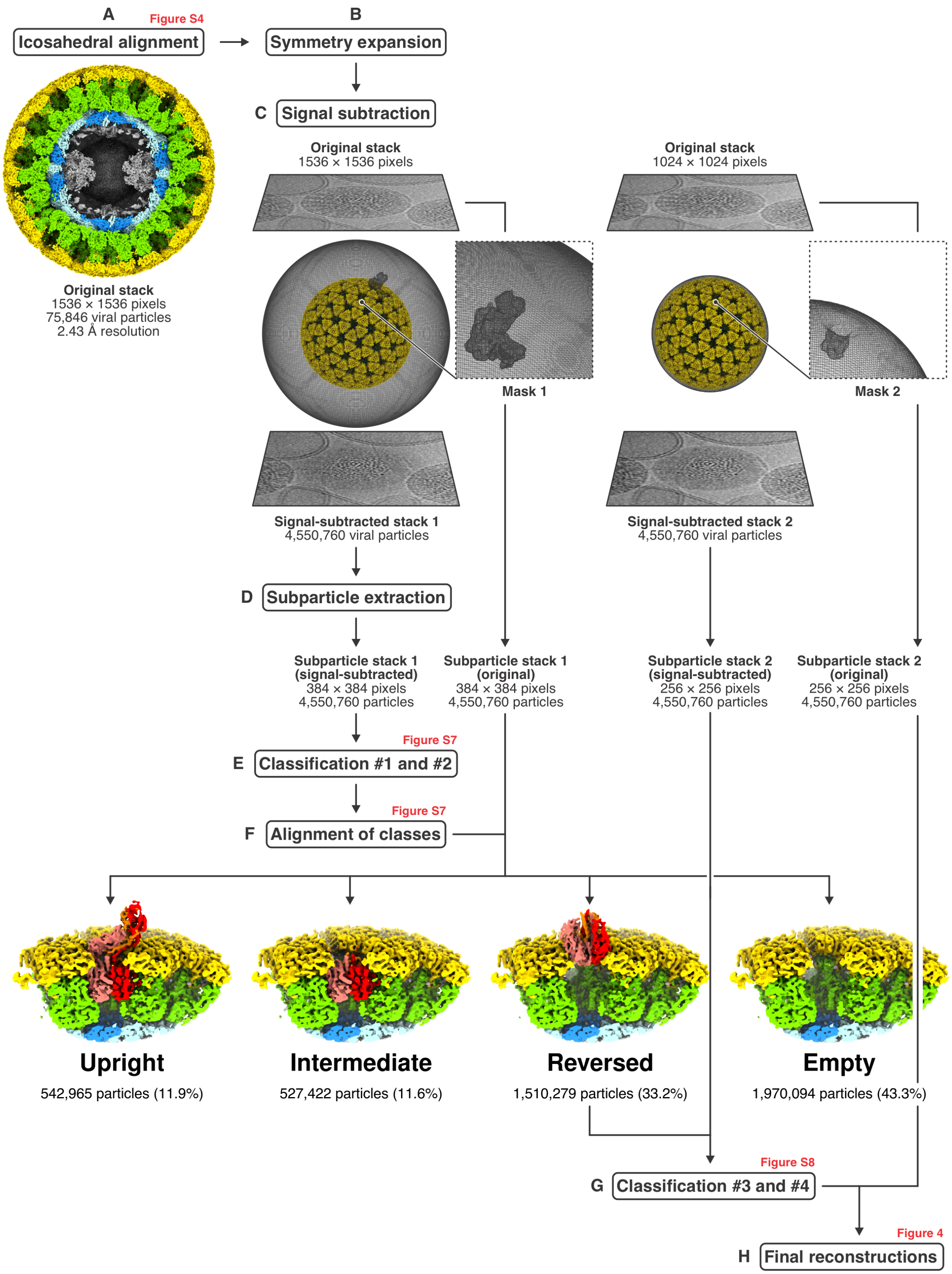


**Figure S5. Cryo-EM analysis processing scheme of TLPs with liposomes.**

(A) Alignment of full viral particles with icosahedral symmetry imposed. A map of the final icosahedral reconstruction is shown, in which the particle is partially cut. VP1 (RNA-dependent

RNA polymerase) and RNAs are colored gray; VP2, cyan and blue; VP6, green; VP7, yellow. See also Figure S4. (B) Icosahedral symmetry expansion (60-fold) based on the full particle alignment. (C) Signal subtraction. The masks used to the define the subtraction volume, excluding the region of interest at a single protomer position, are shown. Signal-subtracted stack 1 was used to prepare subparticles for classifications #1 and #2 (spike conformation and occupancy). Signal-subtracted stack 2 was used to prepare subparticles for classifications #3 and #4 (liposome interaction of reversed spikes). (D) Subparticle extraction from the signal-subtracted and original particles stacks. (E) Classifications #1 and #2, see Figure S7. (F) Alignment of classes to correct for observed shifts in reconstructions of subparticle classes caused primarily by anisotropic magnification distortion, see Figure S6. Subparticle reconstructions are shown for the three observed spike conformations (upright, intermediate, and reversed) and empty positions. Density maps were low pass filtered at 5 Å resolution and partially cut. VP2, cyan and blue; VP6, green; VP7, yellow; VP5*, red, orange, and salmon. (G) Classifications #3 and #4, see Figure S8. (H) Final reconstructions of liposome-bound classes calculated from subparticle stack 2, which was obtained from the original (non-signal-subtracted) images.

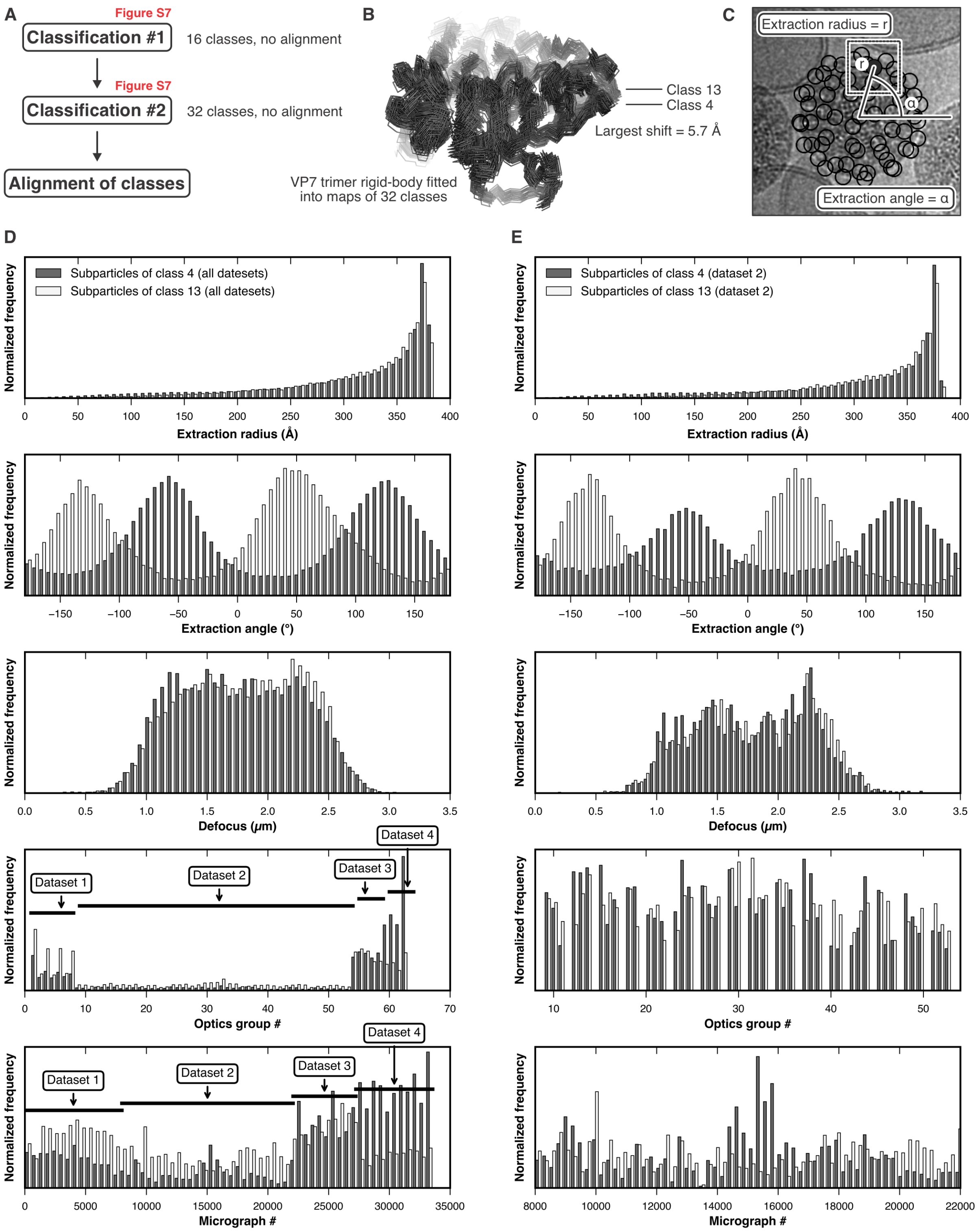


**Figure S6. Observed subparticle shifts caused by anisotropic magnification distortion.**

(A) Subparticles were classified without alignment. Relative shifts between classes were determined by rigid-body fitting a V7 trimer model and then aligning the classes by updating the subparticle alignment parameters based on the fitted models. (B) Ribbon representation of the VP7 trimer models after fitting it into the densities of the 32 classes. The largest shift we observed was 5.7 Å between classes 4 and 13. (C) Each subparticle is associated with an extraction radius and angle with respect to the center of the full virus particle projection in the original micrograph. (D) Histograms of metadata values (extraction radius, extraction angle, defocus, optics group number, micrograph number) for all subparticles from the two classes 4 and 13, which showed the largest relative shift in their reconstructions. A strong bias in the extraction angle suggests that the observed shifts are predominately caused by anisotropic magnification distortion, which was not accounted for during subparticle extraction. (D) Histograms of metadata values (extraction radius, extraction angle, defocus, optics group number, micrograph number) for dataset 2 subparticles from the two classes 4 and 13.

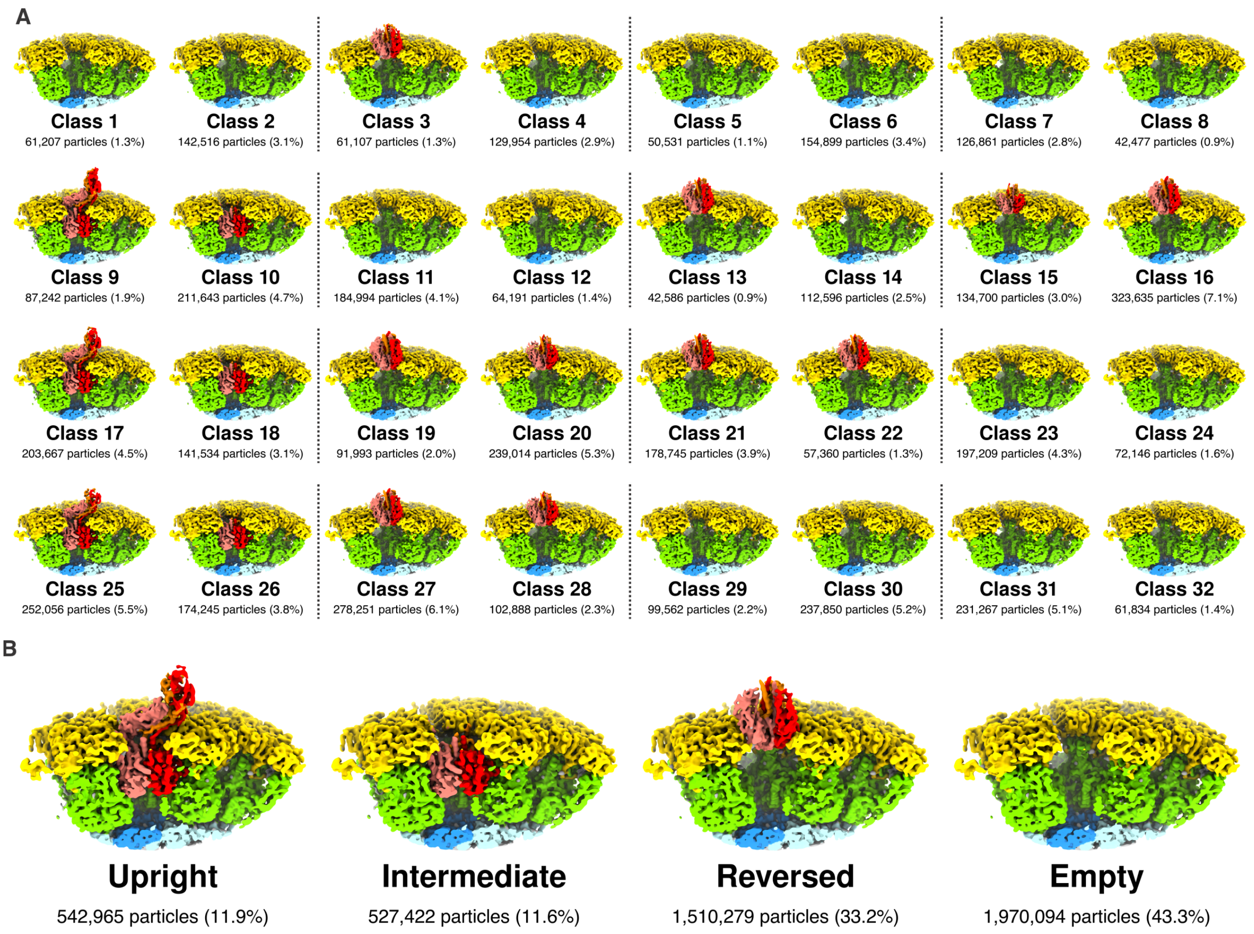


**Figure S7. Classification #1 and #2.**

For display, density maps were calculated from subparticle stack 1, which was obtained from the original (non-signal-subtracted) images, low pass filtered at 5 Å resolution and partially cut. VP2, cyan and blue; VP6, green; VP7, yellow; VP5*, red, orange, and salmon. (A) Classification #1 partitioned the particles into 16 classes, each of which was further subclassified into two classes in classification #2 (indicated by dashed lines). (B) Reconstructions of three observed spike conformations (upright, intermediate, and reversed) and empty positions after merging corresponding classes (Table S2) and correcting of observed subparticle shifts (Figure S6).

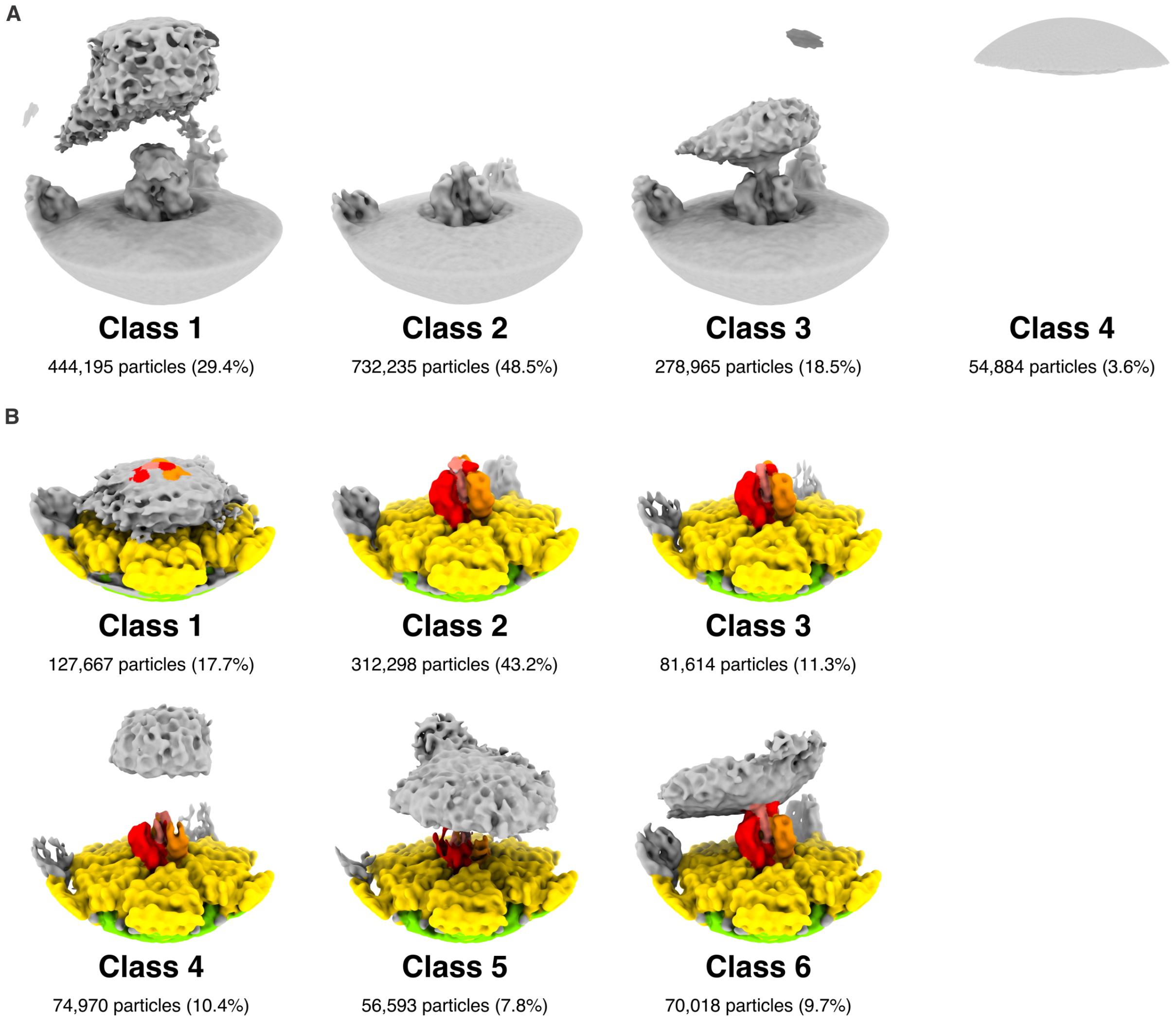


**Figure S8. Classification #3 and #4.**

Classification of liposome-bound spike positions from subparticles with reversed spike conformations. (A) In classification #3, we initially requested four classes. Density maps were calculated from signal-subtracted subparticle stack 2, low pass filtered at 8 Å resolution and displayed in gray and at very low contour level to visualize membrane density. (B) In classification #4, we further subclassified classes 1 and 3 from classification #3. Density maps of the six classes were calculated from subparticle stack 2, which was obtained from the original (non-signal-subtracted) images, and low pass filtered at 8 Å. VP6, green; VP7, yellow; VP5*, red, orange, and salmon; membrane, gray.

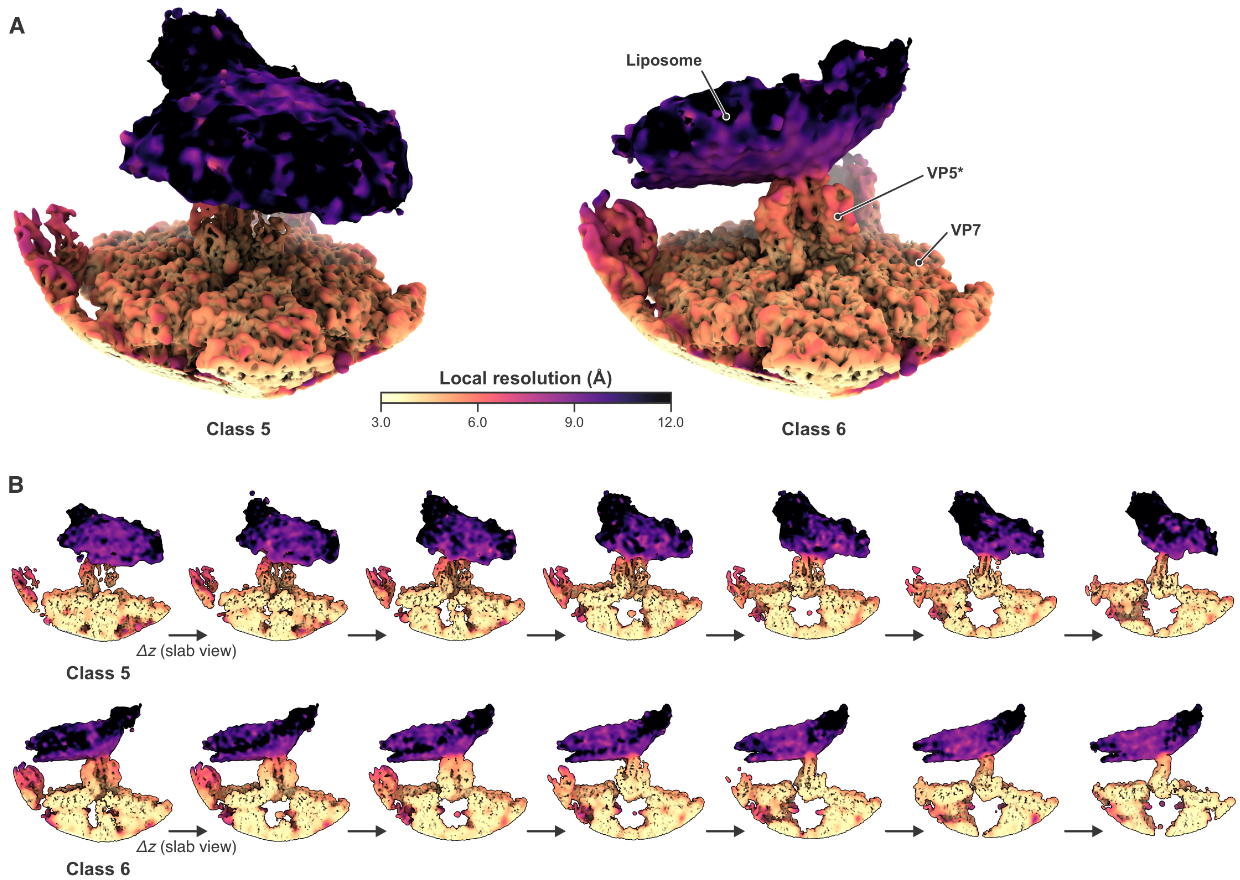


**Figure S9. Local resolution analysis of liposome-bound VP5*.**

(A) Full view of the class 5 and class 6 reconstructions colored according to local resolution. (B) Corresponding slab views.

**Table S1. Cryo-EM data collection and statistics**

| | Dataset 1<br>RRV liposomes | Dataset 2<br>RRV liposomes | Dataset 3<br>RRV liposomes | Dataset 4<br>RRV liposomes |
|---|---|---|---|---|
| **Data collection** | | | | |
| Electron microscope | Titan Krios | Titan Krios | Titan Krios | Titan Krios |
| Camera | K3 Summit | K3 Summit | K3 Summit | K3 Summit |
| Magnification | 60,606 | 60,606 | 60,606 | 60,606 |
| Voltage (kV) | 300 | 300 | 300 | 300 |
| Number of movies | 7,985 | 14,088 | 5,170 | 6,320 |
| Defocus range (μm) ∗ | 1.0–2.5 | 1.0–2.5 | 1.0–2.5 | 1.0–2.5 |
| Pixel size (Å) | 0.825 | 0.825 | 0.825 | 0.825 |
| **Icosahedral reconstruction** | | | | |
| Number of images | 75,846 | | | |
| Box size (pixels) | 1536 | | | |
| Pixel size (Å) | 0.825 | | | |
| Symmetry imposed | I (setting I2) | | | |
| Map resolution (Å) † | 2.43 | | | |
| | **Subparticle stack 1** | **Subparticle stack 2** | **Final class 5**<br>EMD-42343<br>PDB-ID 8UK2 | **Final class 6**<br>EMD-42344<br>PDB-ID 8UK3 |
| **Local reconstruction** | | | | |
| Number of images | 4,550,760 | 4,550,760 | 56,593 | 70,018 |
| Box size (pixels) | 385 | 256 | 256 | 256 |
| Pixel size (Å) | 0.825 | 1.2375 | 1.2375 | 1.2375 |
| Symmetry imposed | $C_1$ | $C_1$ | $C_1$ | $C_1$ |
| Map resolution (Å) | 2.91, 2.73 ‡ | 3.33, 3.14 § | N.A. | N.A. |

∗ Approximate range of underfocus.
† Resolution where Fourier shell correlation (FSC) between half-maps drops below 0.143 after applying a spherical shell mask (inner radius = 222 Å, outer radius = 403 Å).
‡ Resolution after applying a mask encompassing one VP7 trimer and the volume corresponding to VP4 in upright and reversed conformation for reconstructions from the original (non-signal-subtracted) subparticle stack 1 before and after alignment of classes 1–32, respectively.
§ Resolution after applying a mask encompassing volume corresponding to VP4 in upright and reversed conformation for reconstructions from the original (non-signal-subtracted) subparticle stack 2 before and after alignment of classes 1–32, respectively.

**Table S2. Classification #1 and #2 of VP5*/VP8* spike positions**

| Classification #1 * | | | Classification #2 † | | | | |
|---|---|---|---|---|---|---|---|
| 1 | 203,723 | 4.5% | 1 | 1 | 61207 | 1.3% | Empty (partially occupied) |
| | | | 2 | 2 | 142516 | 3.1% | Empty |
| 2 | 191,061 | 4.2% | 1 | 3 | 61107 | 1.3% | Reversed |
| | | | 2 | 4 | 129954 | 2.9% | Empty |
| 3 | 205,430 | 4.5% | 1 | 5 | 50531 | 1.1% | Empty |
| | | | 2 | 6 | 154899 | 3.4% | Empty |
| 4 | 169,338 | 3.7% | 1 | 7 | 126861 | 2.8% | Empty |
| | | | 2 | 8 | 42477 | 0.9% | Empty |
| 5 | 298,885 | 6.6% | 1 | 9 | 87242 | 1.9% | Upright |
| | | | 2 | 10 | 211643 | 4.7% | Intermediate (β-barrel domains disordered) |
| 6 | 249,185 | 5.5% | 1 | 11 | 184994 | 4.1% | Empty |
| | | | 2 | 12 | 64191 | 1.4% | Empty |
| 7 | 155,182 | 3.4% | 1 | 13 | 42586 | 0.9% | Reversed |
| | | | 2 | 14 | 112596 | 2.5% | Empty |
| 8 | 458,335 | 10.1% | 1 | 15 | 134700 | 3.0% | Reversed |
| | | | 2 | 16 | 323635 | 7.1% | Reversed |
| 9 | 345,201 | 7.6% | 1 | 17 | 203667 | 4.5% | Upright |
| | | | 2 | 18 | 141534 | 3.1% | Upright (β-barrel domains disordered) |
| 10 | 331,007 | 7.3% | 1 | 19 | 91993 | 2.0% | Reversed |
| | | | 2 | 20 | 239014 | 5.3% | Reversed |
| 11 | 236,105 | 5.2% | 1 | 21 | 178745 | 3.9% | Reversed |
| | | | 2 | 22 | 57360 | 1.3% | Reversed |
| 12 | 269,355 | 5.9% | 1 | 23 | 197209 | 4.3% | Empty |
| | | | 2 | 24 | 72146 | 1.6% | Empty (partially occupied) |
| 13 | 426,301 | 9.4% | 1 | 25 | 252056 | 5.5% | Upright |
| | | | 2 | 26 | 174245 | 3.8% | Intermediate (β-barrel domains disordered) |
| 14 | 381,139 | 8.4% | 1 | 27 | 278251 | 6.1% | Reversed |
| | | | 2 | 28 | 102888 | 2.3% | Reversed |
| 15 | 337,412 | 7.4% | 1 | 29 | 99562 | 2.2% | Empty |
| | | | 2 | 30 | 237850 | 5.2% | Empty |
| 16 | 293,101 | 6.4% | 1 | 31 | 231267 | 5.1% | Empty |
| | | | 2 | 32 | 61834 | 1.4% | Empty |
| | | | | | **542,965** | **11.9%** | **Upright** |
| | | | | | **527,422** | **11.6%** | **Intermediate** |
| | | | | | **1,510,279** | **33.2%** | **Reversed** |
| | | | | | **1,970,094** | **43.3%** | **Empty** |
| | | | | | **4,550,760** | **100%** | **Total** |

∗ Columns are class number of classification #1, number of particles, percentage of particles.

† Columns are class number of classification #2, final class number, number of particles, percentage of particles, occupancy state and conformation.

**Table S3. Classification #3 and #4 of reversed VP5*/VP8* spikes**

| Classification #3 * | | | |
|---|---|---|---|
| 1 | 444,195 | 29.4% | Liposome density |
| 2 | 732,235 | 48.5% | No liposome density |
| 3 | 278,965 | 18.5% | Liposome density |
| 4 | 54,884 | 3.6% | Junk |
| | **1,510,279** | **100%** | **Total** |

| Classification #4 * | | | |
|---|---|---|---|
| 1 | 127,667 | 17.7% | Liposome density |
| 2 | 312,298 | 43.2% | No liposome density |
| 3 | 81,614 | 11.3% | No liposome density |
| 4 | 74,970 | 10.4% | Liposome density |
| 5 | 56,593 | 7.8% | Liposome density |
| 6 | 70,018 | 9.7% | Liposome density |
| | **723,160** | **100%** | **Total** |

∗ Columns are class number, number of particles, percentage of particles, assignment.
