## Supplementary figures and images for "The rotavirus VP5*/VP8* conformational transition permeabilizes membranes to Ca^2+^"

### Supplemental animation

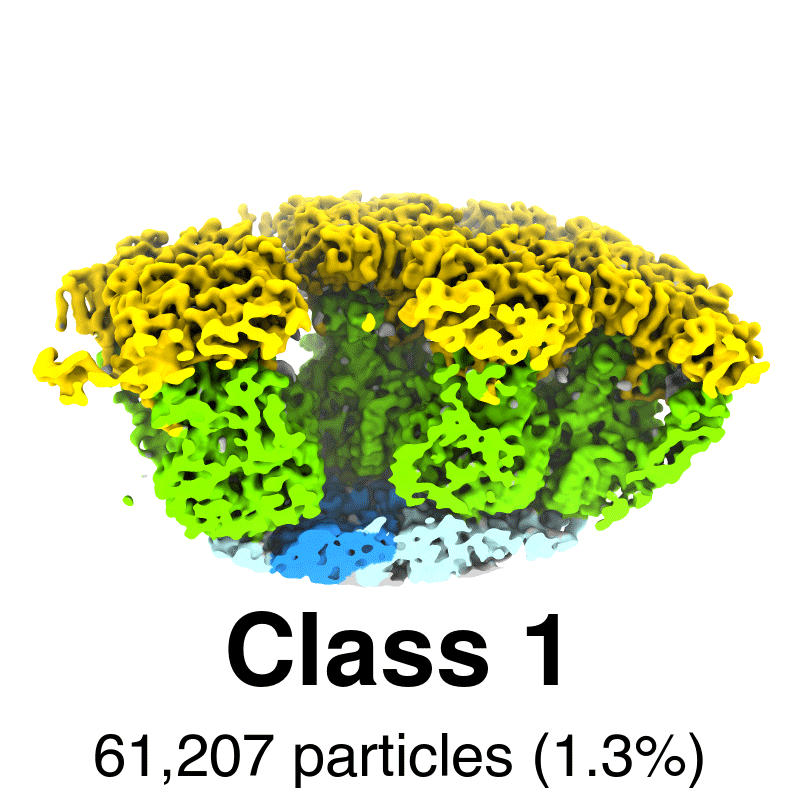
